# Native receptor-targeted chemogenetics enables cell-type-specific inhibition of endogenous receptors in freely moving mice

**DOI:** 10.64898/2026.02.23.707339

**Authors:** Tomohiro Doura, Wataru Kakegawa, Keigo Morikawa, Tomoteru Yamasaki, Duy Phuoc Tran, Masayuki Fujinaga, Takumi Kondo, Shuntaro Kashiwa, Kanta Hasegawa, Eriko Miura, Saeko Matsudaira, Yuya Fujihara, Hiroshi Nonaka, Itaru Hamachi, Michisuke Yuzaki, Akio Kitao, Ming-Rong Zhang, Shigeki Kiyonaka

## Abstract

Understanding brain function requires tools that allow precise manipulation of receptor signaling in defined cell types within intact neural circuits. Optogenetics and conventional chemogenetic approaches primarily enable cell-type-specific control of neuronal excitability using engineered receptors or ion channels. However, direct and reversible inhibition of endogenous neurotransmitter receptors in defined cell types has remained technically inaccessible. Here, we introduce native receptor-targeted chemogenetics (NARCH), a chemogenetic strategy that integrates structure-guided receptor engineering with allosteric ligand design to achieve reversible and temporally precise inhibition of endogenous receptor signaling with cell-type specificity *in vivo*. By applying NARCH to metabotropic glutamate receptor 1 (mGlu1), we demonstrate that mGlu1 signaling in cerebellar Purkinje cells is required for stabilization of motor learning across training sessions in freely moving mice. NARCH thus establishes a receptor-level chemogenetic framework for causal analysis of neural circuits and behavior.

---

Cell-surface receptors serve as fundamental molecular interfaces that convert extracellular signals into intracellular signaling programs, thereby shaping complex cellular and physiological responses. In the brain, neurotransmitter receptors constitute a central molecular basis for highly ordered functions such as learning and memory^1,2^. A defining feature of these receptors is their organization into receptor families composed of multiple subtypes encoded by distinct genes^3–5^. Although each family member is activated by the same endogenous ligand, receptor subtypes exhibit distinct expression patterns across brain regions and contribute to region-specific physiological functions. Thus, elucidating the causal roles of individual receptor subtypes requires approaches that enable selective suppression of endogenous receptor signaling within defined brain regions, rather than global perturbation of receptor activity.

Loss-of-function approaches in mouse models have been widely employed to investigate receptor function *in vivo* (Table 1). Conditional knockout of a receptor gene enables brain region- or cell-type-selective deletion of target receptors and has provided important insights into their roles in learning and memory^6^. However, genetic deletion from early developmental stages often causes severe phenotypes that preclude behavioral analyses, complicating the distinction between acute receptor functions and developmental effects or compensatory adaptations^7,8^. Inducible genetic strategies, such as Tet-on/off systems, offer reversible suppression of receptor expression, but their reliance on transcriptional and translational regulation operates on a timescale of days, limiting temporal resolution for analyzing acute receptor function^9^. By contrast, pharmacological approaches using receptor antagonists offer reversible inhibition with high temporal resolution^10^. However, systemically administered compounds diffuse broadly throughout the brain, limiting cell- or region-specific suppression, and observed phenotypes may be confounded by off-target effects^11,12^.

**Table 1.** Comparison of *in vivo* strategies for receptor manipulation.

|  | Cell-type-specificity<sup>a</sup> | Native receptor targeting | Temporal resolution | Reversibility | Off-target effect<sup>b</sup> |
| --- | --- | --- | --- | --- | --- |
| Genetic deletion | + | + | – | – | + |
| Pharmacology | – | + | + | + | – |
| Conventional chemogenetics | + | – | + | + | + / – |
| Native receptor-targeted chemogenetics (NARCH) | + | + | + | + | + / – |

<sup>a</sup> Cell-type-specificity in genetic deletion refers to conditional knockout strategies.
<sup>b</sup> ‘+’ indicates minimal or negligible off-target effects (*i.e.*, favorable), whereas ‘–’ indicates reported or potential off-target effects.

Chemogenetic approaches that combine genetic engineering with pharmacological control enable reversible and cell-type-specific manipulation of neuronal activity *in vivo*^13,14^. Among these, designer receptors exclusively activated by designer drugs (DREADDs) have been widely used to interrogate brain function by decoupling engineered receptors from endogenous ligand signaling^15^. Despite their utility, conventional chemogenetic approaches primarily modulate neuronal activity through engineered receptors, thereby operating at the level of whole-cell excitability rather than enabling receptor-level inhibition of endogenous signaling.

Here, we introduce <u>na</u>tive <u>r</u>eceptor-targeted <u>ch</u>emogenetics (NARCH), a chemogenetic strategy that enables reversible and cell-type-selective inhibition of endogenous receptor signaling *in vivo* (Fig. 1a and Table 1). As a proof of concept, we focused on metabotropic glutamate receptor subtype 1 (mGlu1), a G protein-coupled receptor highly expressed in discrete brain regions including the cerebellum, thalamus, and olfactory bulb^16,17^. In mice, genetic deletion of the mGlu1 gene (*Grm1*) results in severe motor dysfunction, precluding behavioral analyses of higher-order functions such as motor learning^18^. By contrast, NARCH–mGlu1 enables non-invasive, reversible, and cell-type-specific inhibition of endogenous mGlu1 signaling *in vivo*, thereby allowing functional dissection of mGlu1 within intact neural circuits.

**Fig. 1.**
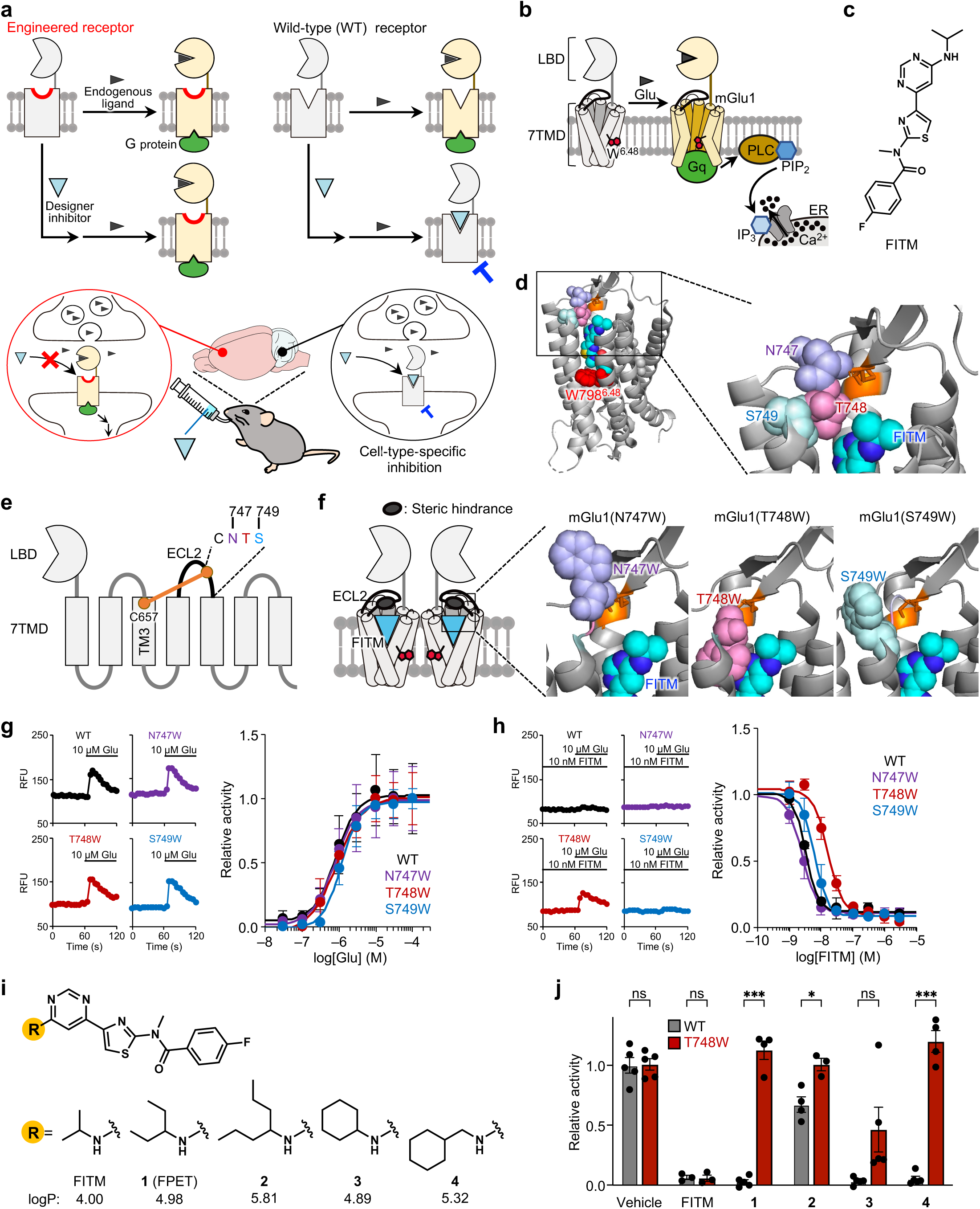
Design and functional screening of FITM derivatives that selectively inhibit WT mGlu1 but not mGlu1(T748W) mutant. **a**, Schematic illustration of NARCH. WT and engineered receptors respond similarly to the endogenous ligand, whereas only the WT is inhibited by the designer inhibitor (upper). This enables cell-type-specific inhibition of endogenous receptor signaling in freely moving mice (lower). **b**, Schematic overview of mGlu1 signaling. Glu, glutamate; PLC, phospholipase C; PIP_2_, phosphatidylinositol 4,5-bisphosphate; IP_3_, inositol 1,4,5-triphosphate; ER, endoplasmic reticulum. **c**, Chemical structure of FITM. **d**, The X-ray crystal structure of the mGlu1 7TMD bound to FITM (PDB ID: 4OR2). Disulfide bond between C657 and C746 is shown in orange. **e**, Schematic presentation of mGlu1. **f**, Schematic presentation of mGlu1 mutants bound to FITM. N747W, T748W, and S749W in ECL2 are shown as medium purple, orchid, and aquamarine spheres, respectively. **g**, Representative Cal-520 fluorescence traces in HEK293 cells expressing WT or mutant mGlu1 (N747W, T748W, and S749W) stimulated by 10 µM Glu (left) and corresponding Glu concentration–response curves (right). Relative activity was calculated as the ratio of the obtained response intensity to that of mGlu1-expressing HEK293 cells induced by 100 µM Glu. Data are presented as mean ± SEM (*N* = 3). **h**, Representative Cal-520 fluorescence traces in HEK293 cells expressing WT or mutant mGlu1 (N747W, T748W, and S749W) pretreated with 10 nM FITM and then stimulated by 10 µM Glu (left) and corresponding FITM concentration–inhibition curves (right). Relative activity was calculated as the ratio of the obtained response intensity to that of mGlu1-expressing HEK293 cells induced by 10 µM Glu in the presence of vehicle. Data are presented as mean ± SEM (*N* = 3). **i**, Chemical structures and calculated logP values of FITM derivatives. **j**, Screening of FITM derivatives for selective inhibition of WT mGlu1 relative to mGlu1(T748W). Relative activity was calculated as the ratio of the obtained response intensity to that of WT mGlu1-expressing HEK293 cells induced by 10 µM Glu in the presence of vehicle. Data are presented as mean ± SEM (*N* = 3–5). \*\*\**p* < 0.001; \*\**p* < 0.01; \**p* < 0.05; ns, not significant. Statistical analyses were performed using the unpaired *t*-test. Amino acid abbreviations: C, Cys; N, Asn; T, Thr; S, Ser; W, Trp.

## Results

### Engineering a chemogenetic mGlu1–inhibitor pair for NARCH–mGlu1

To establish a chemogenetic strategy for selective and reversible inhibition of mGlu1, we developed a designer mGlu1–inhibitor pair in which a designer inhibitor selectively suppresses wild-type (WT) mGlu1 activity while having minimal effect on the engineered receptor (Fig. 1a). Crucially, this strategy requires that the engineered receptor fully retains native receptor function, including normal glutamate responsiveness. mGlu1 is a class C Gq-coupled receptor whose activity is regulated by glutamate binding to the extracellular ligand-binding domain (LBD) as well as allosteric ligands targeting the seven-transmembrane domain (7TMD) (Fig. 1b). We focused on 4-fluoro-*N*-[4-[6-(isopropylamino)pyrimidin-4-yl]-1,3-thiazol-2-yl]-*N*-methylbenzamide (FITM)^19^, an mGlu1-selective negative allosteric modulator (NAM) (Fig. 1c), that binds to 7TMD and stabilizes the inactive state of mGlu1 (Extended Data Fig. 1a–c)^20^. Structural analysis revealed that the 2-propylamino group of FITM is positioned near residues N747–S749 within the extracellular loop 2 (ECL2), a structurally constrained region stabilized by a disulfide bond (Fig. 1d,e and Extended Data Fig. 1d). We therefore hypothesized that introducing mutations into ECL2 residues 747–749 (ECL2^747–749^) would selectively modulate FITM binding, yielding mGlu1 mutants resistant to FITM inhibition. Because the ECL2 is spatially separated from the orthosteric ligand-binding domain and the transmembrane core, mutagenesis in ECL2^747–749^ was expected to preserve glutamate-induced receptor activation.

We constructed mGlu1(N747W), mGlu1(T748W), and mGlu1(S749W) mutants by substituting each residue in ECL2^747–749^ with Trp, a bulky amino acid, to introduce steric hindrance (Fig. 1f). We assessed the activity of each mGlu1 mutant expressed in HEK293 cells via the Ca^2+^ mobilization assay (Supplementary Fig. 1a). All three mGlu1 mutants exhibited glutamate-induced Ca^2+^ responses with concentration dependence comparable to that of WT mGlu1, indicating preserved receptor function (Fig. 1g, Supplementary Fig. 1b, and Supplementary Table 1). Pretreatment with FITM suppressed glutamate-induced activation of each mGlu1 mutant in a concentration-dependent manner (Fig. 1h, Supplementary Fig. 2, and Supplementary Table 2). mGlu1(N747W) and mGlu1(S749W) were inhibited by FITM as well as wild-type (WT) mGlu1. By contrast, mGlu1(T748W) exhibited reduced sensitivity to FITM, with the half-maximal inhibitory concentration (IC_50_) approximately 5-fold higher than that of WT mGlu1 (17 ± 3.3 and 3.3 ± 0.3 nM for mGlu1(T748W) and WT, respectively).

Although mGlu1(T748W) showed reduced sensitivity to FITM, the difference in IC_50_ values between mGlu1(T748W) and WT mGlu1 was considered insufficient for robust *in vivo* applications. To further enhance selectivity, we designed and synthesized FITM derivatives **1**–**4** bearing bulkier substituents than the 2-propylamino group (Fig. 1i). When HEK293 cells expressing WT mGlu1 or mGlu1(T748W) were treated with 3 µM of **1**–**4**, inhibitory potency of **1**, **2**, and **4** against mGlu1(T748W) was markedly reduced (Fig. 1j and Supplementary Fig. 3). Compound **2** also showed reduced inhibition of WT mGlu1, whereas **1** and **4** retained potent inhibition of WT mGlu1. Based on its close structural similarity to FITM and a calculated logP value reflecting hydrophobicity closer to that of FITM, we selected **1** as a designer inhibitor candidate and named it FPET.

### FPET acts as a silent allosteric ligand on mGlu1(T748W)

We then examined the inhibitory action of FPET against WT mGlu1 and mGlu1(T748W) in detail. In HEK293 cells, FPET inhibited glutamate-induced responses of WT mGlu1 in a concentration-dependent manner (IC_50_ = 29 ± 6.1 nM), while FPET failed to inhibit mGlu1(T748W) even at 3 µM (∼100-fold higher than the IC_50_ for WT mGlu1) (Fig. 2a and Supplementary Fig. 4).

**Fig. 2.**
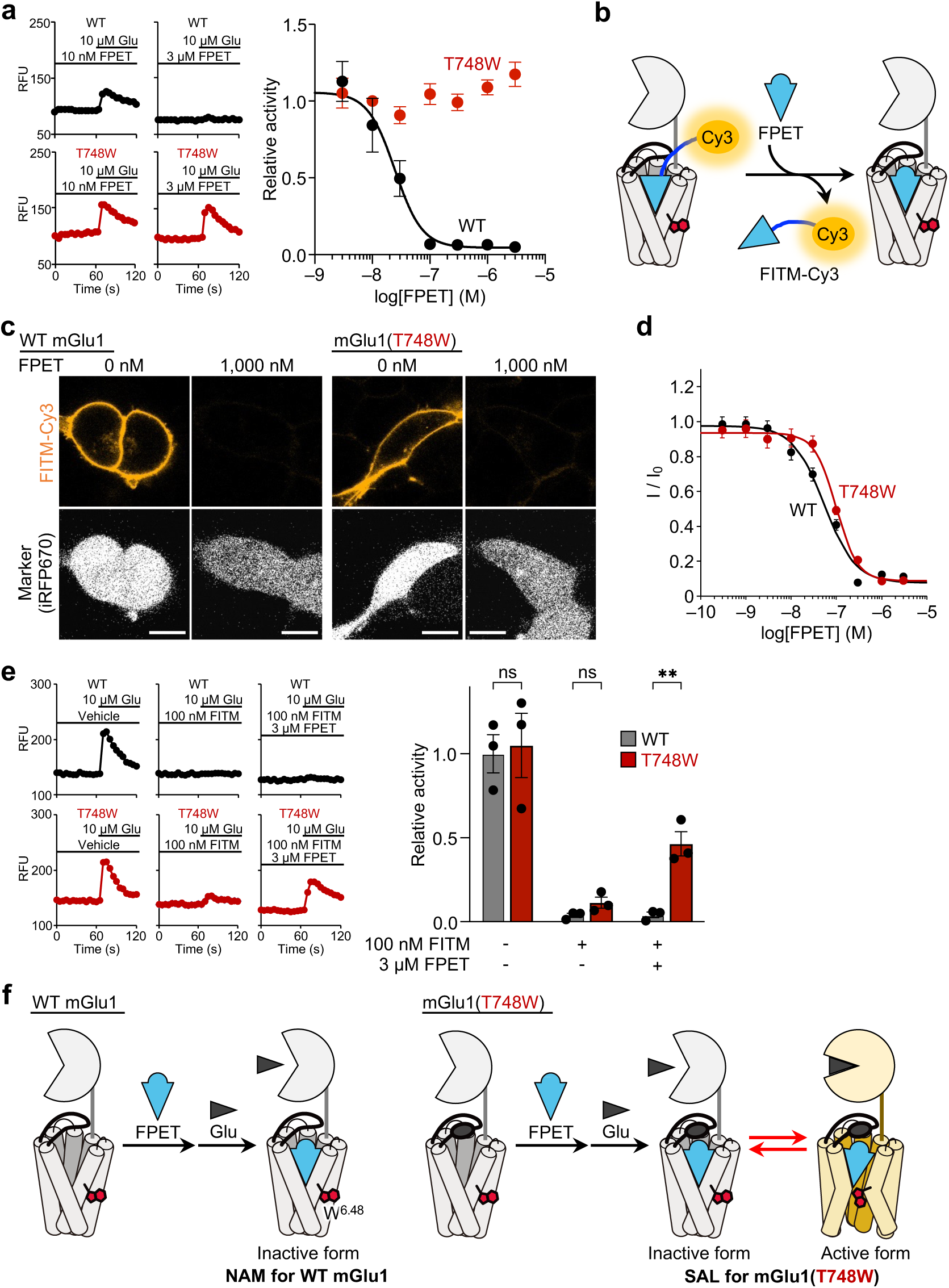
FPET differentially modulates WT mGlu1 and mGlu1(T748W), acting as a NAM or a silent allosteric ligand (SAL), respectively. **a**, Representative Cal-520 fluorescence traces in HEK293 cells expressing WT mGlu1 or mGlu1(T748W) pretreated with FPET (10 nM or 3 µM) then stimulated with 10 µM Glu (left) and corresponding FPET concentration–inhibition curves (right). Relative activity was calculated as the ratio of the obtained response intensity to that of mGlu1-expressing HEK293 cells induced by 10 µM Glu in the presence of vehicle. Data are presented as mean ± SEM (*N* = 3). The IC_50_ of FPET for WT mGlu1 was 29 ± 6.1 nM. **b**, Schematic illustration of competitive binding of FITM-Cy3 and FPET to mGlu1. **c**, Competitive binding assay of FPET in HEK293 cells expressing WT mGlu1 or mGlu1(T748W) using FITM-Cy3 (30 nM for WT mGlu1 and 300 nM for mGlu1(T748W)). iRFP670 was used as a transfection marker. Scale bars, 10 µm. **d**, Concentration-dependent displacement of FITM-Cy3 in HEK293 cells expressing WT mGlu1 or mGlu1(T748W) (*n* = 105 cells). Relative intensity (I/I_0_) was calculated by dividing the obtained fluorescence intensity on the surface of mGlu1-expressing HEK293 cells (I) by that in the presence of vehicle (I_0_). *K*_i_ values of FPET were 9.6 ± 0.7 nM for WT mGlu1 and 35 ± 7.9 nM for mGlu1(T748W). Data are presented as mean ± SEM (*N* = 3). **e**, Representative Cal-520 fluorescence traces in HEK293 cells expressing WT mGlu1 or mGlu1(T748W) treated with vehicle, FITM (100 nM), or FITM (100 nM) plus FPET (3 µM), then stimulated with 10 µM Glu (left) and quantification of the Ca^2+^ responses (right). Relative activity was calculated as the ratio of the obtained response intensity to that of WT mGlu1-expressing HEK293 cells induced by 10 µM Glu in the presence of vehicle. Data are presented as mean ± SEM (*N* = 3). \*\**p* < 0.01; ns, not significant. Statistical analyses were performed using the unpaired *t*-test. **f**, Schematic model illustrating the distinct allosteric effects of FPET on WT mGlu1 and mGlu1(T748W).

The lack of inhibition of mGlu1(T748W) by FPET could arise from either reduced binding affinity or altered allosteric efficacy. To distinguish between these possibilities, we examined FPET binding to mGlu1(T748W) using a competitive assay with FITM-Cy3 (ref. 21), a fluorescent probe for mGlu1 (Fig. 2b and Extended Data Fig. 2a). Confocal imaging showed binding of FITM-Cy3 to both WT and mGlu1(T748W) expressed at the cell surface of HEK293 cells (Extended Data Fig. 2b, c). Importantly, the fluorescence of FITM-Cy3 was markedly decreased upon addition of a high concentration of FPET to cells expressing either WT mGlu1 or mGlu1(T748W) (Fig. 2c and Extended Data Fig. 3). Consistent with this observation, the inhibitory constants (*K*_i_) of FPET were 9.6 ± 0.7 nM for WT mGlu1 and 35 ± 7.9 nM for mGlu1(T748W) (Fig. 2d), indicating that FPET retains substantial binding affinity for the mutant receptor.

We next examined whether FPET alters the allosteric effect on mGlu1. As shown in Fig. 2e, glutamate-induced Ca^2+^ responses mediated by WT mGlu1 and mGlu1(T748W) were effectively suppressed by 100 nM FITM. By contrast, robust Ca^2+^ responses were observed for mGlu1(T748W) when both 100 nM FITM and 3 µM FPET were co-administered, whereas WT mGlu1 was suppressed under the same conditions. These results indicate that FPET competitively binds to mGlu1(T748W) against FITM but lacks negative allosteric efficacy. Consistent with this interpretation, FPET had little effect on the glutamate concentration-response relationship of mGlu1(T748W) (Extended Data Fig. 4). Collectively, FPET functions as a negative allosteric modulator (NAM) on WT mGlu1, but acts as a silent allosteric ligand (SAL) on mGlu1(T748W), binding the receptor without altering its activity (Fig. 2f).

To further examine the molecular basis of this observation, we performed molecular dynamics (MD) simulations of WT mGlu1–FITM and mGlu1(T748W)–FPET complexes. We focused on W798 (W^6.48^) in transmembrane helix 6, a conserved toggle-switch residue whose side-chain orientation is closely associated with GPCR activation^22^. In the X-ray structure of WT mGlu1 in complex with FITM^20^, W^6.48^ is positioned in proximity to the fluorophenyl group of FITM. We conducted ten independent 1 µs MD simulations of the WT mGlu1–FITM complex, which represents an inactive-state conformation stabilized by FITM. Analysis of the χ1 dihedral angle of W^6.48^ (defined by the N–Cα–Cβ–Cγ atoms) revealed that the –170° rotamer was predominantly populated, whereas another rotamer around –80° was weakly populated in both chains A and B (Extended Data Fig. 5). This suggests that the –170° rotamer corresponds to the inactive-like conformation. In contrast, in the ten 1 µs MD simulations of the mGlu1(T748W)–FPET complex, the population of the –80° rotamer was markedly increased. Notably, a similar increase in the –80° population was observed in the simulations of WT mGlu1 in the absence of ligand (apo WT). Given that ligand-free GPCRs exist in equilibrium between inactive and active forms (Extended Data Fig. 1a), the increased –80° population likely corresponds to an active-like conformation. These computational analyses suggest that the mGlu1(T748W)–FPET complex retains a higher propensity to sample active-like conformations, supporting the interpretation that FPET binds to mGlu1(T748W) as a SAL.

### Ligand optimization and *in vivo* pharmacokinetic characterization

Although FPET is suitable for chemogenetic regulation of mGlu1, its calculated logP (4.98) is higher than that of FITM (4.00), a compound with favorable *in vivo* pharmacokinetics (Fig. 1i)^23^. Therefore, we sought FPET analogs with physicochemical properties more similar to those of FITM. Guided by previous structure-activity relationship studies of FITM derivatives^24,25^, we designed and synthesized **5** (CPET) and **6** (MPET) as FPET analogs (Fig. 3a). Among these compounds, MPET exhibited the highest aqueous solubility, consistent with its lower calculated logP value (4.69) (Fig. 3b and Supplementary Fig. 5). Importantly, MPET retained the desired chemogenetic profile, acting as a NAM on WT mGlu1 and as a SAL on mGlu1(T748W) in cell-based assays (Fig. 3c and Supplementary Fig. 6).

**Fig. 3.**
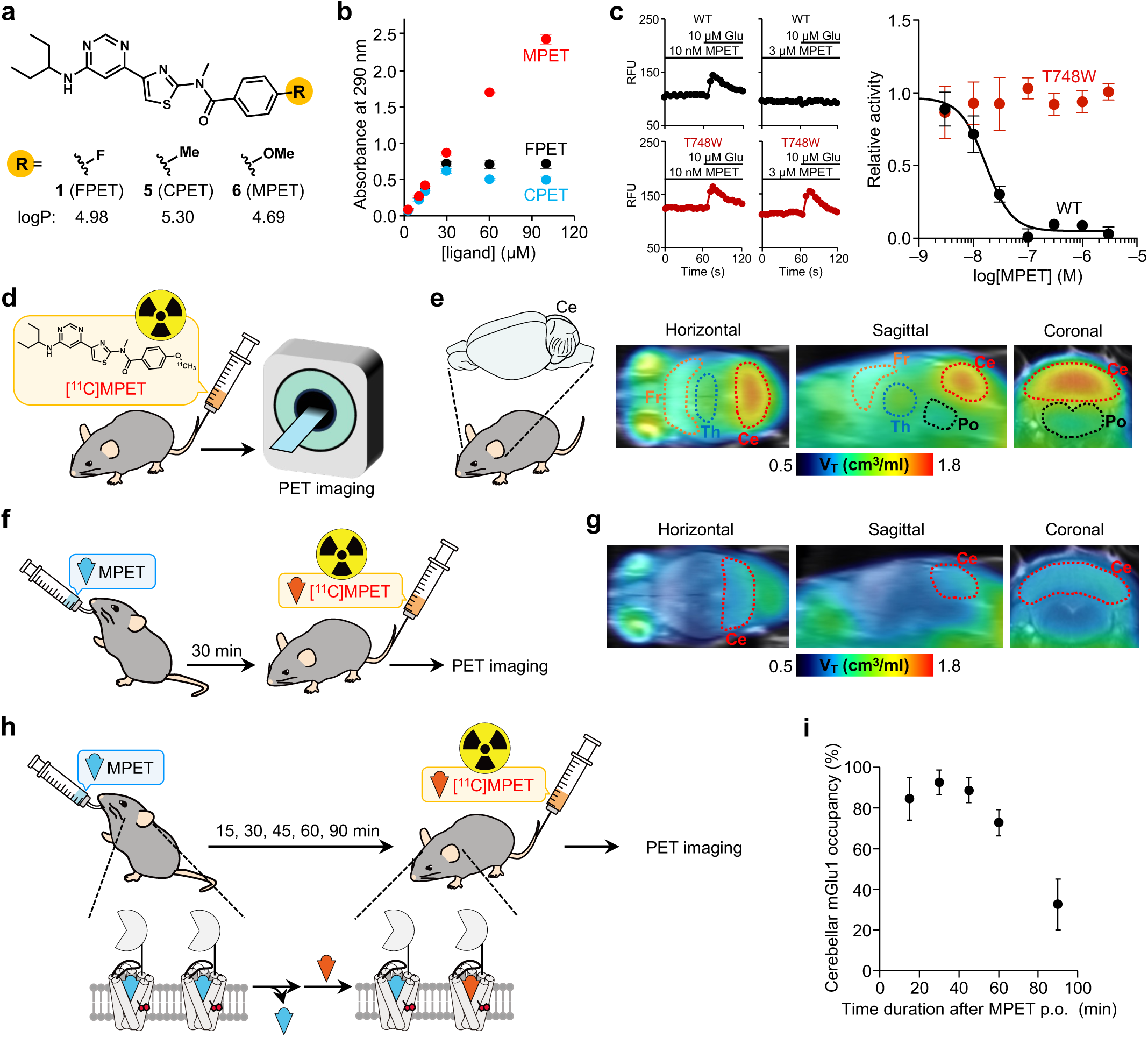
*In vivo* pharmacokinetics and PET-based occupancy analysis of MPET. **a**, Chemical structures and calculated logP values of FPET, CPET, and MPET. **b**, Comparison of aqueous solubility of FPET, CPET, and MPET assessed by absorbance at 290 nm. **c**, Representative Cal-520 fluorescence traces in HEK293 cells expressing WT mGlu1 or mGlu1(T748W) pretreated with MPET (10 nM or 3 µM) then stimulated with 10 µM Glu (left) and corresponding concentration–inhibition curves (right). Relative activity was calculated as the ratio of the obtained response intensity to that of mGlu1-expressing HEK293 cells induced by 10 µM Glu in the presence of vehicle. Data are presented as mean ± SEM (*N* = 3). The IC_50_ of MPET for WT mGlu1 was 17 ± 1.5 nM. **d**, Schematic overview of PET imaging in mice intravenously injected with [^11^C]MPET via tail vein. **e**, Representative PET images of a WT mouse brain (Fr: frontal cortex; Th: thalamus; Ce: cerebellum; Po: pons), co-registered to an MRI template, following intravenous injection of [^11^C]MPET. Dynamic images were reconstructed and scaled by distribution volume (V_T_), which reflects the *in vivo* distribution of [^11^C]MPET with mGlu1. **f**, Schematic overview of PET imaging experiments in mice intravenously injected with [^11^C]MPET after oral administration of MPET. **g**, Representative PET/MRI images of a WT mouse brain following [^11^C]MPET injection 30 min after oral administration of MPET (6 mg/kg). PET images were scaled by V_T_. **h**, Schematic illustration of the experimental design for time-dependent mGlu1 occupancy studies by MPET. **i**, Time-dependent occupancy of mGlu1 by MPET in the cerebellum of WT mice. [^11^C]MPET was injected at 15, 30, 45, 60, or 90 min after oral administration of MPET (6 mg/kg). Data are presented as mean ± SEM (*n* = 3–4 mice per group).

Next, we evaluated the brain penetration of MPET using positron emission tomography (PET). Following intravenous administration of [^11^C]MPET via the tail vein in mice, radioactivity was readily detected throughout the brain and accumulated predominantly in the cerebellum, a region with the highest mGlu1 expression in the brain (Fig. 3d,e and Extended Data Fig. 7a,b)^16,17^. These results indicated that intravenously administered [^11^C]MPET crosses the blood-brain barrier (BBB) efficiently and binds to endogenous mGlu1 *in vivo*.

To assess whether MPET can be administered non-invasively, we next examined whether orally administered MPET reaches the brain and engages endogenous mGlu1. Oral administration of non-radioactive MPET (6 mg/kg) markedly reduced cerebellar radioactivity derived from subsequently administered [^11^C]MPET, indicating that orally administered MPET reaches the brain and occupies mGlu1, thereby preventing binding of [^11^C]MPET to mGlu1 (Fig. 3f,g). PET analyses with [^11^C]MPET at multiple time points after MPET administration further revealed that receptor occupancy by MPET remained at approximately 90% for up to 45 min after oral administration, with a decline evident after 60 min (Fig. 3h,i and Extended Data Fig. 7c,d). These findings indicate that oral administration of MPET enables robust yet reversible inhibition of WT mGlu1 *in vivo* within a defined temporal window.

### Generation and validation of conditional mGlu1(T748W) knock-in mice

Previous studies have shown that mGlu1 knockout (mGlu1-KO) mice exhibit motor dysfunction, including severe ataxia^17,18^. Notably, restoration of mGlu1a expression specifically in cerebellar Purkinje cells rescues motor function in these mice, indicating that mGlu1 signaling in Purkinje cells is essential for normal motor function^18^. However, the chronic loss of mGlu1 in gene knockout models precludes dissection of the contribution of mGlu1 to higher-order brain functions, including motor coordination, motor memory, and motor learning. To enable reversible control of endogenous mGlu1 in a cell-type-specific manner, we generated conditional knock-in mice harboring the T748W mutation in mGlu1 (hereafter referred to as mGlu1^T748W^ mice) (Fig. 4a and Extended Data Fig. 8a–c). In mGlu1^T748W^ mice, the mutant receptor is expressed throughout the body. By contrast, in mGlu1^T748W^(PC–WT) mice, WT mGlu1 is selectively expressed in cerebellar Purkinje cells through Cre-mediated removal of the floxed insertion sequence containing the mGlu1(T748W) mutation, while mGlu1(T748W) expression is preserved in other cell types (Fig. 4a,b).

**Fig. 4.**
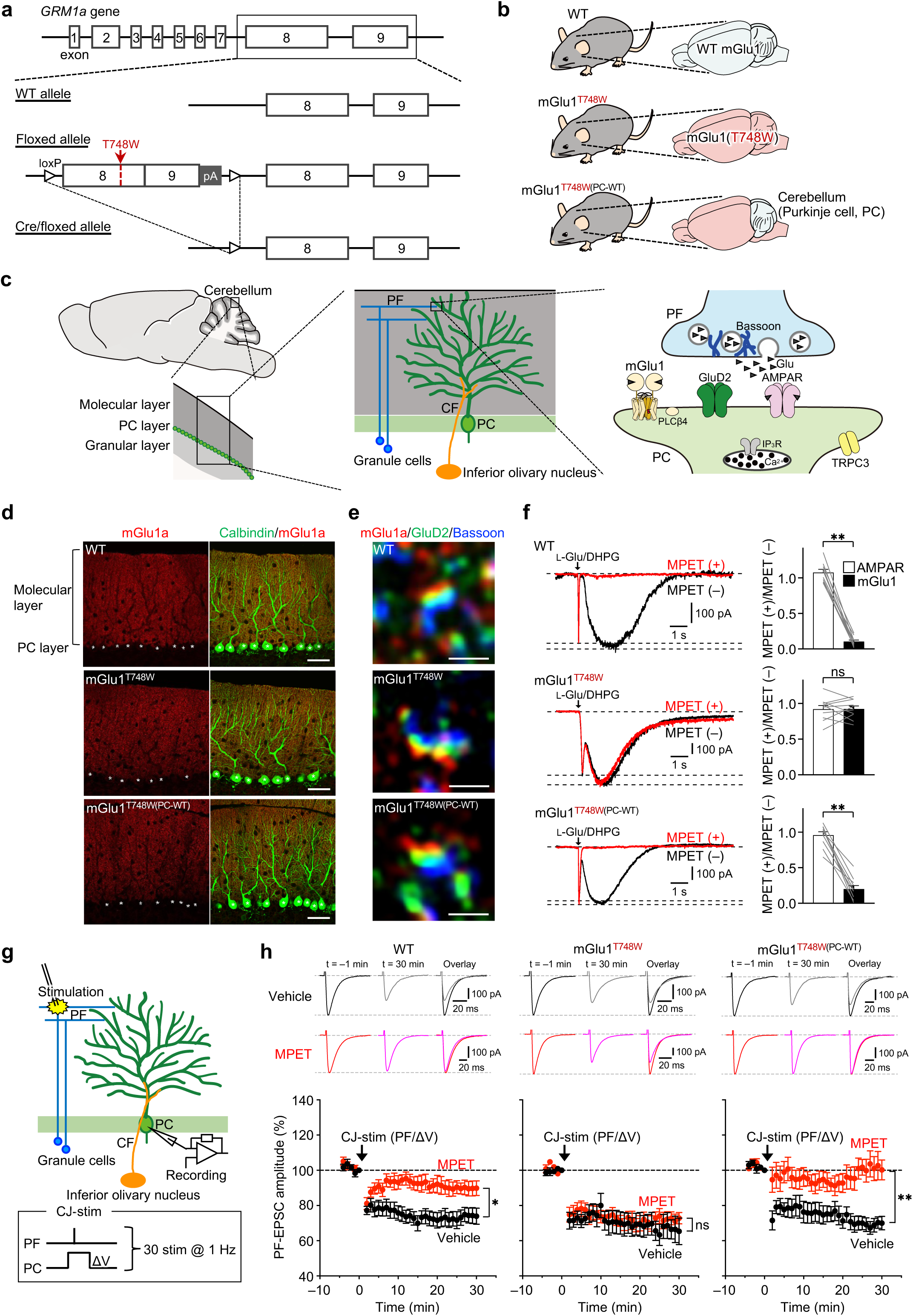
Effects of MPET on endogenous mGlu1 function in the cerebellum. **a**, Strategy for generating mGlu1^T748W^ and mGlu1^T748W(PC-WT)^ mice using the Cre-*loxP* recombination system. **b**, Cartoon representation of the mGlu1^T748W^ and mGlu1^T748W(PC-WT)^ mouse. **c**, Overview of cerebellar organization, showing a sagittal brain section and enlarged view of the cerebellum (left), major components of the cerebellar cortical circuitry (middle), and parallel fiber–Purkinje cell (PF–PC) synapses (right). CF, climbing fiber; AMPAR, AMPA-type glutamate receptor; GluD2, glutamate delta-2 receptor; PLCβ4, phospholipase C β4; IP_3_R, inositol triphosphate receptor; TRPC3, transient receptor potential channel 3. **d**, Immunohistochemical analysis of calbindin (green) and mGlu1a (red) expressed in the molecular layer of the cerebellar cortex from WT, mGlu1^T748W^, and mGlu1^T748W(PC-WT)^ mice. Asterisks indicates Purkinje cell soma. Calbindin is used as a Purkinje cell marker. Scale bars, 50 µm. **e**, Super-resolution images showing immunoreactive mGlu1a (red), GluD2 (green), and Bassoon (blue). GluD2 and Bassoon are used as postsynaptic and presynaptic markers, respectively. Scale bars, 500 nm. **f**, Representative traces with Glu-induced AMPAR currents (fast component) and DHPG-induced mGlu1 currents (slow component). Recordings were performed from the same Purkinje cells under control conditions (black traces) and subsequently after bath application of MPET (red traces). Quantification of AMPAR and mGlu1 currents is shown (right), expressed as the ratio of responses recorded in the presence and absence of MPET for each cell. **g**, Schematic illustration of the experimental paradigm for LTD induction. A conjunctive stimulus [CJ-stim (PF/ΔV), 30 × (PF stimulation paired with Purkinje cell depolarization) at 1 Hz] was applied to induce LTD at PF-Purkinje cell synapses. **h**, Upper, representative PF-EPSC traces immediately before (*t* = –1 min) and 30 min after CJ-stim in WT, mGlu1^T748W^, and mGlu1^T748W(PC-WT)^ mice. Lower, averaged LTD time courses at PF-Purkinje cell synapses. Data are presented as mean ± SEM. \*\**p* < 0.01; \**p* < 0.05; ns, not significant. The data were statistically analyzed using the Wilcoxon signed-rank test in **f** and the Mann–Whitney U test in **h**.

Western blotting analysis revealed comparable expression levels of mGlu1 in the cerebellum of WT, mGlu1^T748W^, and mGlu1^T748W(PC-WT)^ mice (Extended Data Fig. 8d,e). Consistently, immunohistochemical staining using an anti-mGlu1a antibody showed normal mGlu1 distribution in Purkinje cells across these genotypes (Fig. 4c,d). In addition, super-resolution imaging revealed that synaptic localization of mGlu1 was also preserved in these mice (Fig. 4e). Functionally, mGlu1-mediated slow currents evoked by DHPG (mGlu1 currents) showed no significant differences in amplitude among WT, mGlu1^T748W^, and mGlu1^T748W(PC-WT)^ mice (Fig. 4f and Extended Data Fig. 9). In line with these findings, there was no obvious difference in freely moving behavior between WT and mGlu1^T748W^ mice.

### Cell-type-specific inhibition of endogenous mGlu1 in brain slices

We then assessed MPET-mediated inhibition of endogenous mGlu1 function in Purkinje cells using acute cerebellar slice preparations. In cerebellar slices from WT mice, MPET suppressed mGlu1-mediated slow currents without affecting AMPA receptor-mediated fast currents (Fig. 4f). In contrast, MPET failed to inhibit mGlu1 currents in cerebellar slices from mGlu1^T748W^ mice. Importantly, slow mGlu1 currents were suppressed by MPET in cerebellar slices from mGlu1^T748W(PC-WT)^ mice, in which WT mGlu1 is selectively restored in Purkinje cells.

In the cerebellum, Purkinje cells extend numerous highly branched dendrites into the molecular layer, where their distal dendrites form excitatory synapses with parallel fibers (PFs) originating from cerebellar granule cells (Fig. 4c)^26^. Conjunctive stimulation consisting of PF activation and Purkinje cell depolarization (CJ-stim) induces long-term depression (LTD) at PF–Purkinje cell synapses^27^, a form of synaptic plasticity widely considered to underlie motor learning (Fig. 4g). Previous studies have established that mGlu1 activity is essential for the induction of cerebellar LTD^18,28^.

We therefore evaluated the effects of MPET on cerebellar LTD using acute cerebellar slices prepared from WT, mGlu1^T748W^, and mGlu1^T748W(PC-WT)^ mice. In the absence of MPET, CJ-stim induced LTD at PF–Purkinje cell synapses in all three genotypes (Fig. 4h). By contrast, MPET treatment abolished LTD induction in WT and mGlu1^T748W(PC-WT)^ mice, whereas LTD remained intact in mGlu1^T748W^ mice. These results demonstrate that MPET suppresses endogenous WT mGlu1 signaling required for cerebellar LTD, while being functionally inert at the mGlu1(T748W) receptor.

### Chemogenetic inhibition of mGlu1 *in vivo*

We next investigated the cell-type-specific inhibition of mGlu1 *in vivo* using an oculomotor adaptation task that quantitatively assesses cerebellum-dependent sensorimotor adaptation. The horizontal optokinetic response (hOKR) is an ocular reflex that stabilizes retinal images during visual motion, and changes in ocular gain provide a quantitative measure of hOKR performance (Fig. 5a)^29,30^. Consistent with a previous report^31^, mGlu1-KO mice, which exhibit ataxia, failed to show hOKR adaptation (Fig. 5b).

**Fig. 5.**
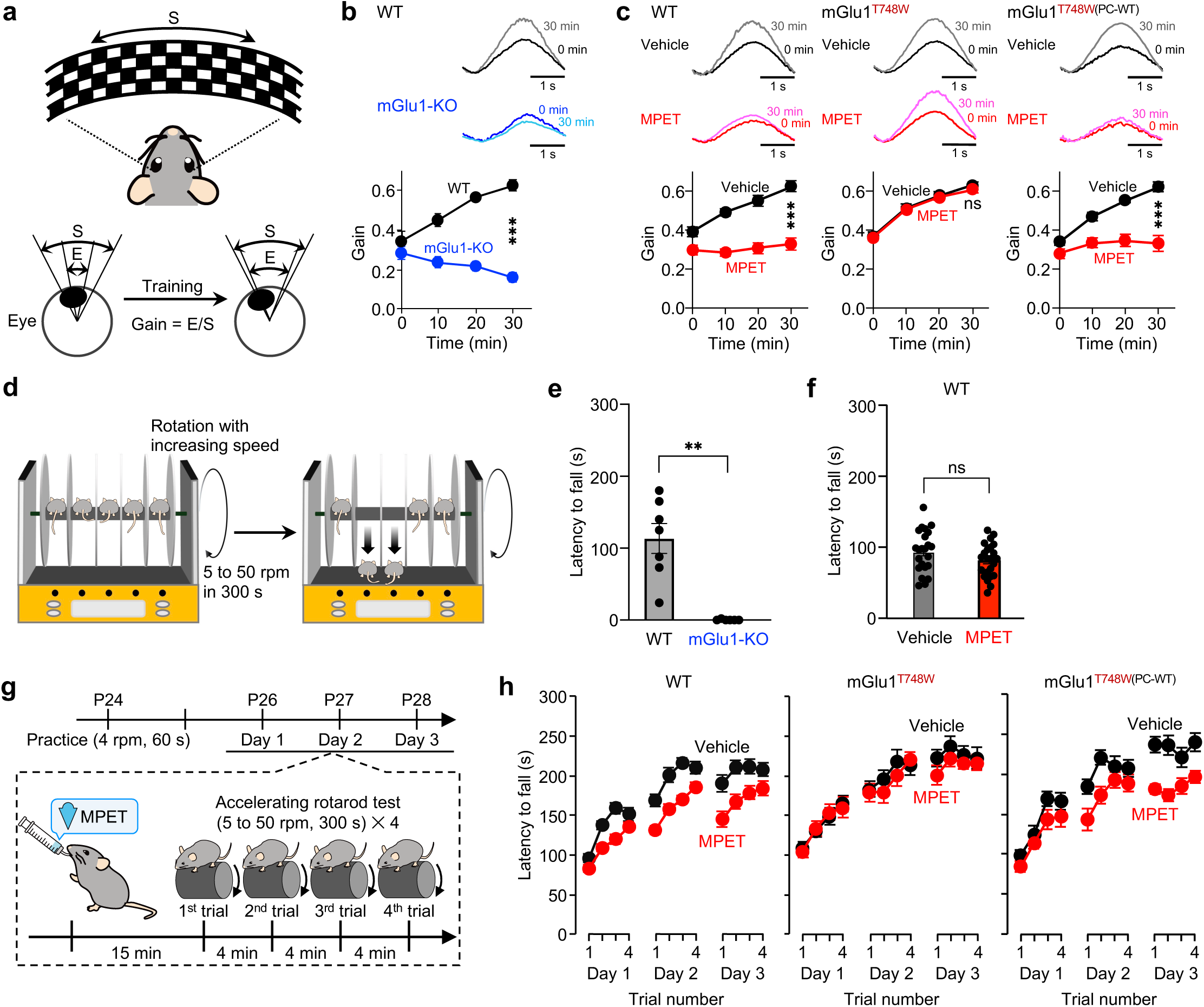
Behavioral analyses reveal critical roles of Purkinje cell-expressed mGlu1 in motor learning. **a–c**, Horizontal optokinetic response (hOKR) adaptation test. In **a**, schematic illustration of the hOKR test is shown. In **b** and **c**, representative hOKR waveforms before training (0 min) and 30 min after the onset of training (upper), and the averaged data of hOKR adaptation (lower) are shown for WT, mGlu1-KO, mGlu1^T748W^, and mGlu1^T748W(PC-WT)^. In **c**, vehicle or MPET (6 mg/kg) was orally administered to each mouse 30 min before the hOKR test. Data are presented as mean ± SEM (WT: *n* = 5 vehicle, 9 MPET; mGlu1^T748W^: *n* = 10 vehicle, 13 MPET; mGlu1^T748W(PC-WT)^: *n* = 7 vehicle, 10 MPET). **d–h**, Accelerating rotarod performance and motor learning. In **d**, schematic overview of the accelerating rotarod test is shown. In **e**, latency to fall for WT and mGlu1-KO mice is shown. Data are presented as mean ± SEM (WT: *n* = 7; mGlu1-KO: *n* = 6). In **f**, latency to fall for WT mice treated with vehicle or 6 mg/kg MPET is shown. Data are presented as mean ± SEM (*n* = 23 per group). In **g**, experimental paradigm for assessing motor learning using the accelerating rotarod test is shown. In **h**, repeated accelerating rotarod test performance over three consecutive days (days 1–3) in WT, mGlu1^T748W^, and mGlu1^T748W(PC-WT)^ mice. Data are presented as mean ± SEM (WT: *n* = 17–21 vehicle, 18–23 MPET; mGlu1^T748W^: *n* = 10–13 vehicle, 11–14 MPET; mGlu1^T748W(PC-WT)^: *n* = 14–17 vehicle, 11–12 MPET). \*\*\**p* < 0.001; \*\**p* < 0.01; ns, not significant. The data were statistically analyzed using two-way repeated measures ANOVA in **b** and **c**, and the unpaired *t*-test in **e** and **f**.

We then examined the effects of orally administered MPET on hOKR performance. Guided by PET-based mGlu1 occupancy measurements (Fig. 3i), hOKR tests were conducted 30 min after oral administration of 6 mg/kg MPET to mice, a time point at which most mGlu1 receptors are occupied by MPET. As shown in Fig. 5c, hOKR adaptation, as measured by ocular gain, was almost completely abolished in MPET-treated WT mice, consistent with effective inhibition of endogenous mGlu1 *in vivo*. By contrast, MPET-treated mGlu1^T748W^ mice exhibited robust hOKR adaptation, with ocular gain comparable to that of vehicle-treated mGlu1^T748W^ mice. This finding excludes non-specific effects of MPET on hOKR adaptation. Importantly, MPET-treated WT mice recovered to vehicle-treated levels 1 day after MPET administration (Extended Data Fig. 10a,b), indicating that MPET inhibition is reversible *in vivo*.

In mGlu1^T748W(PC-WT)^ mice, in which WT mGlu1 is selectively restored in Purkinje cells, MPET suppressed hOKR adaptation to a similar extent as in WT mice (Fig. 5c, right). Because mGlu1(T748W) remains expressed in all other cell types, this result indicates that chemogenetic inhibition of mGlu1 specifically in Purkinje cells is sufficient to impair hOKR adaptation *in vivo*. These findings demonstrate that NARCH enables cell-type-specific and reversible control of endogenous mGlu1 function at the behavioral level.

### Cerebellar mGlu1 is required for motor learning *in vivo*

Complex motor behaviors require both motor coordination and motor learning, which are supported by cerebellar function. However, the acute contribution of cerebellar mGlu1 signaling to these processes remains unclear. To address this, we used the accelerating rotarod test, in which performance within individual trials reflects motor coordination, whereas improvement across repeated trials reflects motor learning (Fig. 5d)^32,33^. In conventional mGlu1-KO mice, severe ataxia precluded reliable assessment of motor learning (Fig. 5e). By contrast, the acute and reversible inhibition of mGlu1 by MPET in the NARCH system enabled evaluation of mGlu1 function in complex motor behaviors *in vivo*.

We first assessed motor coordination in WT mice administered 6 mg/kg MPET or vehicle. MPET did not alter spontaneous locomotor behavior, and retention times on the rotating rod were comparable between MPET-treated and vehicle-treated WT mice (Fig. 5f), indicating that acute mGlu1 inhibition does not substantially impair baseline motor performance.

To assess motor learning, mice were subjected to four rotarod trials per day over three consecutive days (Fig. 5g). The performance during the first trial of day 1 was comparable between MPET-treated and vehicle-treated WT mice, indicating intact baseline coordination. However, across repeated trials, MPET-treated WT mice showed reduced improvement in rotarod performance compared with vehicle-treated controls (Fig. 5h, left). This effect was not observed in mGlu1^T748W^ mice (Fig. 5h, middle), ruling out non-specific effects of MPET.

We further examined the role of mGlu1 in Purkinje cells. In mGlu1^T748W(PC-WT)^ mice, MPET similarly attenuated improvement in rotarod performance across trials (Fig. 5h, right), demonstrating that mGlu1 activity in cerebellar Purkinje cells is required for motor learning. Notably, in mGlu1^T748W(PC-WT)^ mice, MPET did not significantly affect performance at the fourth (final) trial on day 1 but markedly impaired performance at the first trial on day 2 (Fig. 5h, right and Extended Data Fig. 10c). A similar pattern was observed between days 2 and 3, suggesting an impairment in processes contributing to learning consolidation rather than initial acquisition. Together, these results support the conclusion that mGlu1 signaling in cerebellar Purkinje cells contributes to the stabilization of motor learning across training sessions *in vivo*.

## Discussion

In this study, we developed NARCH, a chemogenetic strategy that enables reversible and cell-type-specific control of endogenous receptor function *in vivo*. By applying NARCH to mGlu1, we achieved acute and cell-type-specific inhibition of native mGlu1 signaling without perturbing basal receptor expression or glutamate-induced signaling. Using NARCH-mGlu1, we demonstrate that mGlu1 selectively controls distinct aspects of motor behavior *in vivo*. Acute inhibition of mGlu1 in Purkinje cells impaired motor learning without affecting motor coordination. Furthermore, we show that mGlu1 signaling in Purkinje cells is required for the consolidation, but not necessarily for the acquisition, of motor learning. These findings highlight the power of NARCH to uncover receptor functions with both temporal precision and cell-type specificity that are inaccessible to conventional pharmacological or genetic loss-of-function approaches.

Chemogenetics has emerged as a powerful complement to optogenetics for probing neural circuit function, particularly in contexts where non-invasive modulation of deep brain regions, freely moving animals, or larger animal models including non-human primates is required, owing to its non-invasive mode of action^13,14^. In particular, DREADDs^15^, which employ BBB-permeable designer ligands such as clozapine-*N*-oxide^34^ and deschloroclozapine^35^, have been widely used to bidirectionally modulate neuronal excitability. More recently, a modular chemogenetic platform, PAGER, has been developed in which engineered GPCRs are gated by antigen-dependent relief of auto-inhibition, enabling conditional and programmable control of GPCR signaling at the cellular level^36^. In parallel, chemogenetic control of ion channels has been achieved using the PSAM-PSEM system, which allows selective excitation or inhibition of target neurons through engineered ligand-ion channel pairs^37,38^. Although powerful, these approaches primarily regulate neuronal excitability or output rather than directly manipulating endogenous receptor-mediated signaling pathways, limiting their ability to resolve receptor subtype-specific functions within intact neural circuits.

Several chemogenetic strategies have been developed to achieve protein-level control of receptors, including DART^39^ and DART2^40^. In these approaches, HaloTag proteins^41^ displayed on the surface of target cells are covalently labeled with chimeric ligands composed of a HaloTag-reactive ligand and a receptor-targeting moiety, enabling cell-type-specific regulation of endogenous receptors. However, these chimeric compounds lack BBB-permeability and therefore require intracerebral administration, and covalent tethering of the ligand to HaloTag precludes reversible control of receptor function. Related extensions of this framework, such as maPORTL^42,43^, incorporate additional modalities including photoswitchable ligands^44^, but remain limited by challenges associated with *in vivo* delivery. Recent reports exploring alternative routes toward receptor-selective chemogenetic control further highlight the growing interest in receptor-level manipulation, while differing in mechanism and scope from the inhibition-based strategy described here^45^. In this context, NARCH employs BBB-permeable ligands and minimally perturbed receptors, thereby enabling reversible and temporally precise inhibition of endogenous receptor function *in vivo*.

A key advantage of NARCH is that it enables acute and temporally restricted suppression of endogenous receptor function in the adult brain. This allowed us to directly compare the effects of post-developmental mGlu1 inhibition with those observed in mGlu1 knockout (KO) mice, in which the receptor is deleted from early development. Both acute inhibition of mGlu1 by NARCH and genetic deletion in mGlu1-KO mice produced convergent and divergent effects on cerebellar motor function. In the hOKR paradigm, both mGlu1-KO mice and mice treated with NARCH exhibited impaired performance, consistent with a requirement for mGlu1 signaling in cerebellum-dependent behavior. However, whereas mGlu1-KO mice exhibited severe deficits in the accelerated rotarod test, acute suppression of mGlu1 with NARCH caused no detectable impairment. These findings suggest that coordinated locomotion, as assessed by the rotarod, depends on mGlu1 during developmental circuit formation rather than on ongoing signaling activity in the mature cerebellum. Alternatively, the acquisition of coordinated locomotion may require cumulative limb-based motor experience during development that is, at least in part, mGlu1-dependent. Thus, by enabling temporally restricted and reversible receptor manipulation, NARCH allows the adult contribution of cerebellar mGlu1 to be disentangled from its developmental roles and provides a general framework for dissecting stage-specific receptor functions *in vivo*.

Severe phenotypes caused by constitutive genetic deletion often obscure receptor functions in complex brain processes, making it difficult to distinguish developmental effects from ongoing signaling requirements. By enabling reversible and temporally controlled manipulation of endogenous receptors in defined cell types, NARCH overcomes this limitation of conventional genetic approaches. This capability allows receptor functions to be interrogated at specific time points in the mature brain, independently of developmental or compensatory adaptations. Together, these findings highlight the potential of NARCH as a receptor chemogenetic strategy for resolving stage-specific receptor functions with high temporal and cellular precision.

## Methods

### Synthesis

All synthetic procedures and compound characterizations are described in the Supplementary information.

### Construction of expression vectors

Site-directed mutagenesis was performed using Q5^®^ Site-Directed Mutagenesis Kit (NEB) or NEBuilder HiFi DNA Assembly Master Mix (NEB) with the pCDM vector encoding rat mGlu1^46^ according to the manufacturer’s instructions. The pCDM vector derived from pcDNA3.1(+) (Invitrogen), in which the neomycin cassette was excised using *Pvu*II.

### Cell culture and transfection

HEK293 cells (ATCC) were maintained in Dulbecco’s modified Eagle’s medium (DMEM) (Gibco, Nacalai) supplemented with 100 units/ml penicillin and 100 µg/ml streptomycin and 10% fetal bovine serum (FBS) (Sigma-Aldrich, Nichirei) at 37 °C in a humidified atmosphere of 95% air and 5% CO_2_. HEK293 cells were transfected with an expression plasmid encoding mGlu1 using Lipofectamine 3000 (Thermo Fisher Scientific) according to the manufacturer’s instruction. After transfection, cells were cultured in DMEM GlutaMAX (Gibco) supplemented with 100 units/ml penicillin and 100 µg/ml streptomycin and 10% dialyzed FBS (Gibco, Serana Europe) replacing 10% FBS to decrease cytotoxicity. After 4 h, the medium was removed and fresh DMEM GlutaMAX supplemented with 100 units/ml penicillin and 100 µg/ml streptomycin and 10% dialyzed FBS was added to the transfected cells.

### Ca^2+^ mobilization assay

The transfected HEK293 cells were seeded on a 96-well advanced TC plate (Greiner Bio-One) at 2.0 × 10^4^ cells per well and incubated for 14–16 h at 37 °C under 5% CO_2_. Cells were loaded with 5 µM Cal-520 AM (AAT Bioquest) in DMEM GlutaMAX containing penicillin, streptomycin and dialyzed FBS, and incubated for 4 h at 37 °C under 5% CO_2_. A test ligand was diluted to each prescribed concentration with pre-warmed HEPES-buffered saline (HBS; 20 mM HEPES, 107 mM NaCl, 6 mM KCl, 1.2 mM MgSO_4_, 2 mM CaCl_2_, 11.5 mM D-glucose at pH 7.4) and kept at 37 °C. The cells were washed three times with HBS using an AquaMax 2000 (Molecular Devices). Test ligand was added to the cells, and 3 min later, a mixture of Glu and test ligand was added to stimulate the cells. The fluorescence intensity of Cal-520 (excitation 490 nm, emission 525 nm) per well was monitored using a FlexStation 3 (Molecular Device). The fluorescence increase after stimulation was calculated by subtracting average fluorescence intensity before addition of Glu from the maximum fluorescence intensity induced by Glu. The fluorescence increase (*y*) against Glu concentration (*x*) was fitted with KaleidaGraph (Synergy Software) to calculate the half-maximal effective concentration (EC_50_) value using the following equation:

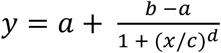

where *a*, *b*, *c* and *d* represent the minimum and maximum fluorescence increase, EC_50_ value, and Hill coefficient, respectively.

### Confocal live cell imaging of cell-surface mGlu1 in HEK293 cells

HEK293 cells were co-transfected with an expression plasmid encoding mGlu1 and piRFP670-N1 (a gift from Dr. V. Verkhusha, Addgene plasmid #45457)^47^ as a transfection marker. Cells were washed with HBS twice, and incubated with FITM-Cy3 (ref. 21) or a mixture of FITM-Cy3 and ligand (FPET or MPET) for 1 h at room temperature in the dark. Confocal live cell imaging was performed with LSM900 (ZEISS) equipped with a 63× oil-immersion objective (NA 1.4). Fluorescence images were acquired with excitation at 561 or 640 nm using diode lasers. To quantify the fluorescence intensity of Cy3 at the mGlu1-expressing cell surface, the maximum fluorescence intensity of the iRFP670-positive cell surface was measured after background subtraction using ZEN Blue software (ZEISS). The plot of Cy3 fluorescence intensity (*y*) against ligand concentration (*x*) was fitted with KaleidaGraph (Synergy Software) to calculate the *K*_d_ value of FITM-Cy3 and the *K*_i_ value of ligand using the following equation:

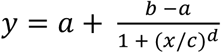

where *a*, *b*, *c* and *d* represent the minimum and maximum Cy3 fluorescence intensity of the cell surface, the half-maximal ligand concentration, and Hill coefficient, respectively.

### Molecular dynamics (MD) simulation

All MD simulations were performed using the AMBER22/24. We employed the X-ray structure of WT mGlu1 in complex with FITM (PDB ID: 4OR2^20^) as the initial structure for the MD simulations of the WT mGlu1–FITM and mGlu1(T748W)–FPET complexes. This structure also served as a template to build the model of the mGlu1(T748W)–FPET complex using HHPred^48^ and Modeller^49^. The structures were then embedded into a membrane consisting of 1-palmitoyl-2-oleoyl-sn-glycero-3-phosphocholine (POPC) and cholesterol (CHL) at a 4:1 ratio to mimic cell membrane conditions. We used the AMBERff19SB^50^, GAFF2^51^, and Lipid21^52^ force fields to model the protein, ligand, and lipids, respectively. The constructed models were then solvated in an OPC^53^ water box with ionic strength of 0.15 M KCl. Li-Merz parameters were applied for ionic interactions^54^. Each system was first energy-minimized by 100,000 steps of steepest descent followed by 100,000 steps of conjugate gradient minimization. The system was heated from 50 K to 300 K over 1 ns MD using the NVT ensemble, followed by 1 ns MD with the NPT ensemble relaxation at 300 K and 1 bar using Langevin dynamics. During this step, positional restraints were applied to the heavy atoms of the ligand, protein backbone and lipid headgroups with a force constant of 100 kcal mol^−1^ Å^−2^. The positional restraints on the lipid headgroups were gradually released over 10 ns. The system was further equilibrated for an additional 10 ns with positional restraints applied solely to the backbone atoms with a force constant of 100 kcal mol^−1^ Å^−2^. Production simulations were carried out without positional restraints for 2 µs. Ten independent 2 µs simulations were performed using the Monte Carlo barostat^55^. For the apo-state simulations, the final snapshots from the ten WT mGlu1–FITM simulations were used to generate ten independent initial configurations by removing the ligands. The resulting systems were subsequently re-equilibrated for 10 ns under strong positional restraints (100 kcal mol^−1^ Å^−2^) applied to all protein atoms, the headgroups of POPC lipids, and the hydroxyl groups of cholesterol. Following re-equilibration, 1 µs production simulations were performed for each system.

### Radiosynthesis of [^11^C]MPET

Carbon-11 (^11^C) was produced by ^14^N (p,α) ^11^C nuclear reactions using CYPRIS HM-18 cyclotron (Sumitomo Heavy Industries). [^11^C]MeI was synthesized from cyclotron-produced [^11^C]CO_2_ as previously described^56^. Briefly, [^11^C]CO_2_ was bubbled into 0.04 M LiAlH_4_ in anhydrous tetrahydrofuran (THF, 300 μl). After evaporation of THF, the remaining complex was treated with 57% hydroiodic acid (300 μl) to give [^11^C]MeI, which was distilled at 180 °C and transferred with N_2_ gas into a solution of **14** (1.0 mg) and NaOH (5 μl, 0.5 M) in anhydrous DMF (300 μl) at –15 to ‒20 °C. After radioactivity reached a plateau, reaction mixture was heated at 70 °C for 5 min. The reaction mixture was injected onto a semipreparative HPLC system. HPLC columns were used as follows: CAPCELL PAK UG80 (10 mm × 250 mm) (Osaka-Soda) for separation; CAPCELL PAK UG80 (4.6 mm × 250 mm) (Osaka-Soda) for analysis. HPLC purification was completed using the mobile phase of MeCN/H_2_O/Et_3_N (65/35/0.01, v/v/v) at a flow rate of 5.0 ml/min. The radioactive fraction corresponding to the desired product was collected in a sterile flask, evaporated to dryness in vacuo, redissolved in 3 ml of sterile normal saline, and passed through a 0.22 μm Millipore filter to give 3.8 GBq of [^11^C]MPET. The retention time of [^11^C]MPET was 8.5 min for purification and 12.7 min for analysis on HPLC with MeCN/H_2_O/Et_3_N (60/40/0.01, v/v/v) at a flow rate of 1.0 ml/min. The total synthesis time from end-of-bombardment was 28 min, providing a decay-corrected radiochemical yield of 27% based on [^11^C]CO_2_, a radiochemical purity of >99%, and a molar activity (*A*_m_) of 90 GBq/μmol at the end of synthesis.

### Animals

All experimental procedures were performed in accordance with the National Institutes of Health Guide for the Care and Use of Laboratory Animals, and were approved by the Institutional Animal Care and Use Committees of Nagoya University, Gakushuin University, Keio University, and National Institutes for Quantum Science and Technology. Wild-type (WT) C57BL/6N mice maintained under specific pathogen-free conditions were purchased from Japan SLC, Inc (Shizuoka, Japan). Animals were housed under controlled environmental conditions (23 ± 1 °C, 12-h light/dark cycle) with ad libitum access to food and water in accordance with the Guidance for Proper Conduct of Animal Experiments by the Ministry of Education, Culture, Sports, Science, and Technology of Japan.

### PET imaging in mice

WT male mice (3–4 weeks old, 14.6 ± 2.0 g) were anesthetized with isoflurane, a catheter was inserted into the tail vein for [^11^C]MPET injection, and the animals were positioned at the center of an Inveon PET scanner (Siemens Healthineers). Following a bolus injection of [^11^C]MPET (9.5 ± 4.0 MBq, 0.1 ml), dynamic PET scans were acquired in 3D list mode for 60 min (1 min × 4 scans, 2 min × 8 scans, 5 min × 8 scans). The acquired data were reconstructed into dynamic PET images using filtered back projection with a Hanning filter (Nyquist cutoff, 0.5 cycle/pixel). Using PMOD software (version 3.4; PMOD Technologies), volumes of interest for mGlu1 were placed on the cerebellum, thalamus, frontal cortex, and pons by referencing MRI templates of a typical C57BL/6J mouse brain, and regional time-activity curves of [^11^C]MPET were extracted. Radioactivity was decay-corrected and expressed as % injected dose per milliliter (%ID/ml). Parametric PET images scaled to distribution volume (*V*_T_) were generated using Logan graphical analysis with an image-derived input function in PMOD^57^.

### Estimating mGlu1 occupancy in the cerebellum

Occupancy of non-radiolabeled MPET at brain mGlu1 receptors was estimated as previously described^58^. In addition to baseline scans, PET experiments were performed in WT mice at multiple time points (15, 30, 45, 60, and 90 min) after oral administration of MPET (6 mg/kg). To determine the non-displaceable distribution volume (*V*_ND_), PET scans under complete receptor blockade were conducted in mice pretreated intravenously with FITM (10 mg/kg), followed by Lassen plot analysis^59^. The cerebellar binding potential (*BP*_ND_), a quantitative dimensionless index of receptor density^60^, was estimated using the following equation:

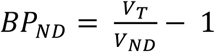

mGlu1 occupancy in the cerebellum was calculated as the percentage reduction in *BP*_ND_
from baseline:

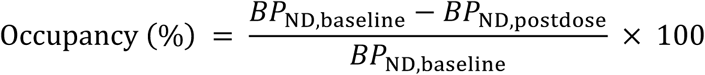

where *BP*_ND,baseline_ and *BP*_ND,postdose_ represent *BP*_ND_ values obtained from PET analyses under baseline conditions and after MPET pretreatment, respectively.

### Generation of mGlu1^T748W^ and mGlu1^T748W(PC-WT)^ mice

As shown in Fig. 4a, the mouse *Grm1* gene encoding mGlu1 comprises nine exons, and the codon for T748 (ACC) is located in exon 8. To generate mGlu1^T748W^ mice, a targeting vector containing a loxP-flanked cassette consisting of the endogenous intron 7 splice acceptor, the T748W mutant coding sequence of exons 8–9, the 3’ untranslated region (UTR) of mouse *Grm1*, an SV40 polyadenylation (pA) signal, and an FRT-flanked neomycin resistance (Neo) cassette was constructed. The *Grm1* targeting construct was linearized by NotI digestion and electroporated into C57BL/6N embryonic stem (ES) cells according to the standard procedures of Cyagen Biosciences. Transfected ES cells were subjected to G418 selection (200 µg/ml) for 24 h after electroporation. G418-resistant clones were isolated and expanded in 96-well plates. After confirmation of correct targeting by PCR screening and Southern blot analysis, a positive ES cell clone was injected into C57BL/6N albino embryos, which were subsequently implanted into CD-1 pseudopregnant females. Founder animals were identified by coat color, and germline transmission was confirmed by crossing with Flp-deleter females followed by genotyping of the F1 offspring. Genotypes were determined by PCR using loxP-specific primers (*forward*, 5’-CTGCACTTGACAGTTTAAGCAATAC-3’; *reverse*, 5’-ATTGTAAGCAGTTGTCACCGC-3’), yielding a 269-bp product for WT alleles and 269- and 325-bp products for flox/+ alleles. T748W mutation (ACC to TGG) was verified by Sanger sequencing of a 358 bp PCR product which was obtained by PCR with the following primers (*forward*, 5’-CAAGAAGAAGATCTGTACCCGGAA-3’; *reverse*, 5’-GCTCCCAAAGTAAATGGGAACAAA-3’). The above procedures to obtain flox/+ mice were outsourced to Cyagen Biosciences. Heterozygous (flox/+) mice were intercrossed to obtain homozygous floxed (flox/flox) mice as shown in Extended Data Fig. 8a. Because T748W mutation introduces an MfeI restriction site into the floxed allele, genotypes were further confirmed by PCR with the following primers (*forward* F1, 5’-CGTAAACGGCCACAAGTTCGAATAAC-3’; *reverse* R1, 5’-CACAGACTTGCCG-TTAGAATTGGCGTTC-3’), yielding a 1,480-bp product, followed by MfeI digestion to give 439-, 475- and 566-bp products for flox/flox mice as shown in Extended Data Fig. 8c. Exon 8 integrity was also verified by PCR with the following primers (*forward* F2, 5’-TTTGTCTATAAGCCTATATGCTTACAGATAAGGGC-3’; *reverse* R2, 5’-GACCT-AGAACCTCAGATTAGACAACTCAC-3’), yielding a 1,368-bp product and MfeI digestion to give 353- and 1,015-bp products for flox/flox mice as shown in Extended Data Fig. 8c.

For generation of mGlu1^T748W(PC-WT)^ mice, mGlu1^T748W^ mice were crossed with Pcp2-Cre mice (The Jackson Laboratory). Presence of the Cre recombinase transgene was confirmed by PCR using Cre-specific primers (*forward*, 5’-GCGGTCTGGCAGT-AAAAACTATC-3’; *reverse*, 5’-GTGAAACAGCATTGCTGTCACTT-3’), yielding a 102-bp product.

### Western blotting analysis of the cerebellum

Cerebella isolated from WT, mGlu1^T748W^, or mGlu1^T748W(PC-WT)^ mice were homogenized in radioimmunoprecipitation assay (RIPA) buffer (50 mM Tris-HCl, 150 mM NaCl, 1 mM EDTA, 1% NP-40, 0.5% sodium deoxycholate, and 0.1% SDS, pH 7.4) containing 1% protease inhibitor cocktail (Nacalai Tesque). Homogenates were sonicated and centrifuged to collect the supernatants. Protein samples were diluted with RIPA buffer containing 1% protease inhibitor cocktail to a final concentration of 0.25 µg/µl. Aliquots of the diluted supernatant (100 µl) were mixed with 25 µl of 5 × Laemmli sample buffer (300 mM Tris-HCl, 15% SDS, 19.7% sucrose, and 0.05% bromophenol blue, pH 6.8) containing 250 mM DTT and incubated for 30 min at room temperature with shaking. Samples were loaded onto SDS-PAGE gels (Bio-Rad Mini-Protean III) and transferred onto polyvinylidene fluoride (PVDF) membranes, followed by blocking with Bullet Blocking One (Nacalai Tesque) for 5 min at room temperature. Membranes were incubated overnight at 4 °C with the following primary antibodies diluted in Tris-buffered saline (10 mM Tris-HCl and 150 mM NaCl, pH 8.0) containing 0.05% Tween 20 (TBS-T) supplemented with 5% Bullet Blocking One: mouse monoclonal anti-mGlu1 (BD Biosciences, 610964, 1:3,000), rabbit monoclonal anti-GluA2 (Abcam, ab206293, 1:3,000), and mouse monoclonal anti-β-actin (MBL, M177-3, 1:3,000). After washing three times with TBS-T, membranes were incubated with HRP-conjugated secondary antibodies (goat polyclonal anti-mouse IgG, MBL, 330, 1:3,000; goat polyclonal anti-rabbit IgG, MBL, 458, 1:3,000) for 1 h at room temperature. After washing three times with TBS-T, chemiluminescent signals generated with ECL Start or ECL Prime (GE Healthcare) were detected using a Fusion Solo S imaging system (Vilber Lourmat). Band intensities of mGlu1 and GluA2 were normalized to the intensity of β-actin.

### Immunohistochemistry of cerebellar slices

Under deep anesthesia with pentobarbital, mice were perfused transcardially with 0.1 M sodium phosphate buffer (PB, pH 7.4) containing 4% paraformaldehyde (4% PFA/PB). Brains were excised and postfixed in 4% PFA/PB for 2 h. After rinsing with PB, parasagittal sections (50 µm thick) were cut using a microslicer (DTK-2000, D.S.K.). Sections were permeabilized and blocked in PB containing 0.1–0.2% Triton X-100 (Sigma-Aldrich) supplemented with either 10% normal donkey serum or 2% normal goat serum and 2% bovine serum albumin (BSA) for 20 min. Sections were incubated overnight at room temperature with primary antibodies: anti-mGlu1α (1:1,000, Frontier Institute), anti-calbindin (1:10,000, Swant), anti-GluD2 (1:500, Frontier Institute), and anti-Bassoon (1:500, Enzo). After washing, sections were incubated for 1 h with appropriate fluorescent dye–conjugated secondary antibodies: anti-rabbit IgG Alexa Fluor 488, anti-guinea pig DyLight 405, and anti-goat IgG Alexa Fluor Plus 647 (1:1,000, Invitrogen). Sections were mounted with Fluoromount-G (SouthernBiotech). Images were acquired using a confocal laser-scanning microscope (FV1000, Olympus) or a super-resolution microscope (OSR, Olympus).

### Preparation of acute cerebellar slices

Mice (4–8 weeks old) were anesthetized with isoflurane, and brains were rapidly removed and transferred to an ice-cold cutting solution containing (in mM): 120 choline chloride, 3 KCl, 1.25 NaH_2_PO_4_, 28 NaHCO_3_, 8 MgCl_2_, 22 D-glucose, and 0.5 sodium ascorbate. The solution was continuously bubbled with 95% O_2_ and 5% CO_2_. Brains were incubated in the cutting solution for 5 min. Sagittal cerebellar slices (200 µm thick) were cut using a microslicer (Linear slicer Pro7, D.S.K.) in ice-cold cutting solution. Slices were then transferred to artificial cerebrospinal fluid (ACSF) containing (in mM):125 NaCl, 2.5 KCl, 2 CaCl_2_, 1 MgCl_2_, 26 NaHCO_3_, 1.25 NaH_2_PO_4_, and 10 D-glucose, bubbled with 95% O_2_ and 5% CO_2_, and incubated at room temperature for at least 1 h before experiments.

### Electrophysiology

Whole-cell patch-clamp recordings were obtained from visually identified Purkinje cells using a 60× water-immersion objective attached to an upright microscope (BX51WI, Olympus) at room temperature, as previously described^61^. Patch pipettes (1.5–3 MΩ) were filled with intracellular solution containing (in mM): 65 Cs-methanesulfonate, 65 K-gluconate, 20 HEPES, 10 KCl, 1 MgCl_2_, 4 Na_2_ATP, 1 Na_2_GTP, 5 sucrose, and 0.4 EGTA (pH 7.25, 295 mOsm/kg) for LTD experiments. For agonist-induced inward current recordings, pipettes were filled with solution containing (in mM): 150 Cs-gluconate, 10 HEPES, 4 MgCl_2_, 4 Na_2_ATP, 1 Na_2_GTP, 0.4 EGTA, and 5 lidocaine *N*-ethyl bromide (QX-314, Tocris Bioscience). Picrotoxin (100 µM; Sigma-Aldrich) was routinely added to ACSF to block inhibitory synaptic transmission. MPET (10 µM) was applied to ACSF during recordings.

To evoke EPSCs derived from PF inputs onto Purkinje cells (PF-EPSCs), square pulses were delivered through a stimulating electrode placed in the molecular layer (50 µm from the pial surface). PF stimulation was confirmed by paired-pulse facilitation at a 50 ms interstimulus interval. For LTD experiments, PF-EPSCs were recorded at a frequency of 0.1 Hz from Purkinje cells voltage-clamped at –80 mV. After stable PF-EPSC amplitudes were observed for at least 5 min, conjunctive stimulation (CJ-stim) consisting of 30 single PF stimuli paired with a 200 ms depolarizing pulse from a holding potential of –60 to +20 mV was applied to induce LTD. Access resistance was monitored every 10 s by measuring peak currents in response to 2 mV, 50 ms hyperpolarizing voltage steps, and recordings were excluded if resistance changed by >20% of its original value. LTD magnitude was calculated from PF-EPSC amplitudes at t = 30 min after CJ-stim.

AMPA receptor-mediated fast currents and subsequent mGlu1-mediated slow currents were evoked by focal perfusion of a mixture of glutamate and DHPG (100 μM each; 10 psi, 50 ms duration) onto dendrites of recorded Purkinje cells (V_h_ = −80 mV). Responses were recorded from the same Purkinje cell under control conditions and then after bath application of MPET (10 μM). AMPA receptor- and mGlu1-mediated currents were quantified as the ratio of post-to pre-application responses for each cell.

Current responses were recorded with an Axopatch 200B amplifier (Molecular Devices), and data were acquired and analyzed with pClamp software (versions 9.2, Molecular Devices). Signals were filtered at 1 kHz and digitized at 4–5 kHz.

### hOKR experiments

Adult mice (at least 6 weeks old) were anesthetized with an intraperitoneal injection of a mixture of medetomidine, midazolam, and butorphanol. A 1 cm flat-head screw was affixed to the skull using dental cement (Super-Bond, Sun Medical). After recovery for at least 1 day, mice were placed on the experimental platform with the head immobilized via the cranial screw and the body loosely restrained in a plastic cylinder. Eye movements were recorded using an infrared LED light source (850 nm LED, Shiokaze Engineering) and a USB 3.0 monochrome camera (DMK23U618, The Imaging Source). Automated pupil tracking was performed with a target-tracking program (Pupil Tracker for OKR-VOR version 6, KATANO TOOL SOFTWARE). Offline analysis using Select Wave software (SW9110, version 3.0.0.1, KATANO TOOL SOFTWARE) removed artifacts caused by saccades and blinks from the acceleration data.

hOKR measurements were conducted using sinusoidal oscillations of a vertically oriented checkered-pattern screen positioned 55 cm from the eye (2 cm square checks) at a frequency of 0.33 Hz, with a peak-to-peak amplitude of 15°, under light conditions (300–400 lux). Eye movement traces free of blinks and saccades were averaged over 10 consecutive cycles, and mean amplitudes were calculated using a modified Fourier analysis, as previously described^62^. To assess hOKR adaptability, mice were exposed to sustained sinusoidal screen oscillation for 30 min. hOKR gain was defined as the ratio of the peak-to-peak amplitude of eye movements to that of the screen oscillation. MPET was administered intragastrically via a cannula at a dose of 6 mg/kg (200 μl) in 0.5% (w/v) methylcellulose 400 (Fujifilm Wako) containing 5–9% dimethyl sulfoxide (DMSO, Fujifilm Wako), or vehicle alone, 30 min before testing after 1 h food and water deprivation.

### Rotarod test

To assess motor coordination in WT and mGlu1-homozygous knockout (–/–) mice^21^, we conducted the accelerating rotarod test as previously described^32^. In brief, littermates obtained by mating mGlu1-heterozygous knockout (+/–) mice were used. At postnatal day 24 (P24), mice were placed on a rotarod treadmill (30 mm diameter; MK-610A, Muromachi Kikai), and acclimated at 4 rotations per minute (rpm) for 60 s. If a mouse could not be placed on the rod after three attempts, acclimation was discontinued. Two days later, mice were tested on a rod accelerating from 5 to 50 rpm for 300 s, and latency to fall was recorded. After testing, genotypes were determined by PCR with the following primers (*forward*, 5’-TGTTTGGCAATACCACCCTC-3’; *reverse*, 5’-TGAGTGAG-AACTCTCCCACG-3’). This produced a 359-bp product for WT mice, whereas a 354-bp product was digested with PvuII to yield 150- and 204-bp fragments for mGlu1 (–/–) mice.

To evaluate motor learning in WT, mGlu1^T748W^, and mGlu1^T748W(PC-WT)^ mice treated with MPET, WT mice were purchased and littermates were obtained by crossing mGlu1^T748W^ and mGlu1^T748W(PC-WT)^ mice. At P24, mice were acclimated on the rotarod at 4 rpm for 60 s as described above. Two days later, MPET (6 mg/kg, 100 µl) in 0.5% (w/v) methylcellulose 400 containing 5–9% DMSO or vehicle alone was administered orally 15 min before testing. Mice were tested on a rotarod accelerating from 5 to 50 rpm for 300 s, and latency to fall was recorded. Each mouse underwent four trials separated by 4-min intervals, and testing was repeated for three consecutive days. After testing, genotypes were confirmed as described in the “Generation of mGlu1^T748W^ and mGlu1^T748W(PC-WT)^ mice” section.

### Statistical analysis

Data are presented as mean ± standard error of the mean (SEM). Values were obtained from at least three independent experiments. Statistical analyses were performed using GraphPad Prism (GraphPad Software) or BellCurve for Excel (Social Survey Research Information). Significance levels were defined as \*\*\**p* < 0.001, \*\**p* < 0.01, and \**p* < 0.05. For comparisons between two groups, an unpaired *t*-test, Mann‒Whitney U test, or Wilcoxon signed-rank test was used as appropriate. For multiple comparisons against a control group, Dunnett’s test was applied. For comparisons among multiple groups, the Kruskal–Wallis test was used.

## Data availability

All data supporting the findings of this study are available within the paper and its Supplementary Information.

## Acknowledgements

The authors acknowledge the Chemical Instrumentation Facility, Nagoya University, for technical assistance and the use of a compact (Bruker) equipped with electron spray ionization (ESI). The authors thank the staff at the National Institutes for Quantum Science and Technology for their assistance with the cyclotron operation, radioisotope production, radiosynthesis, and PET study. We thank Cyagen Biosciences Inc. for generation of mGlu1^T748W^ mice. This work was funded by Grants-in-Aid for Scientific Research (KAKENHI) (Grant Number 22K05351 to T.D., 24K02212, and 22K19364 to W.K., 23K07122 to T.Y., 23K14154 to D.P.T., 22K21353, and 25H01011 to M.Y., 23H02445, 23H02424, 23H04058, 24H01357, and 24H02259 to A.K., 23K27558, and 24K21295 to M.-R.Z., 24H02265, 24H00492, and 24K21823 to S. Kiyonaka), the Naito Foundation (to W.K.), the Brain Science Foundation (to W.K.), the Novartis Foundation (to S. Kiyonaka), the Takeda Science Foundation (to W.K. an S. Kiyonaka) and supported AMED Grant Number 24zf0127012 to S. Kiyonaka

## Author contributions

T.D., W.K. and S. Kiyonaka conceived the project. T.D., K.M., T.K., S. Kashiwa and K.H. designed, synthesized, and characterized compounds. T.D., T.K., S. Kashiwa and K.H. performed the construction of mGlu1 plasmids. T.D., K.M., T.K., S. Kashiwa and K.H. performed cell-based assays. D.P.T., Y.F. and A.K. performed molecular dynamics simulations. T.D., K.M., T.Y., M.F., T.K., M.-R.Z. and S. Kiyonaka designed and performed in vivo PET imaging of mice. T.D., K.M., T.K., S. Kashiwa, K.H. H.N., I.H. and S. Kiyonaka designed and analyzed mGlu1-KO, mGlu1^T748W^ and mGlu1^T748W(PC-WT)^ mice. W.K. and M.Y. designed and performed electrophysiological recordings. W.K. and E.M. performed immunohistocemistory. T.D., W.K. K.M., T.K., S. Kashiwa, K.H., S.M. and S. Kiyonaka designed and performed behavioral tests of mice. T.D., W.K., K.M., M.Y. and S. Kiyonaka wrote the manuscript with input and edits from other authors. W.K., A.K., M.-R.Z. and S. Kiyonaka supervised the manuscript.

---

**Extended Data Fig. 1.**
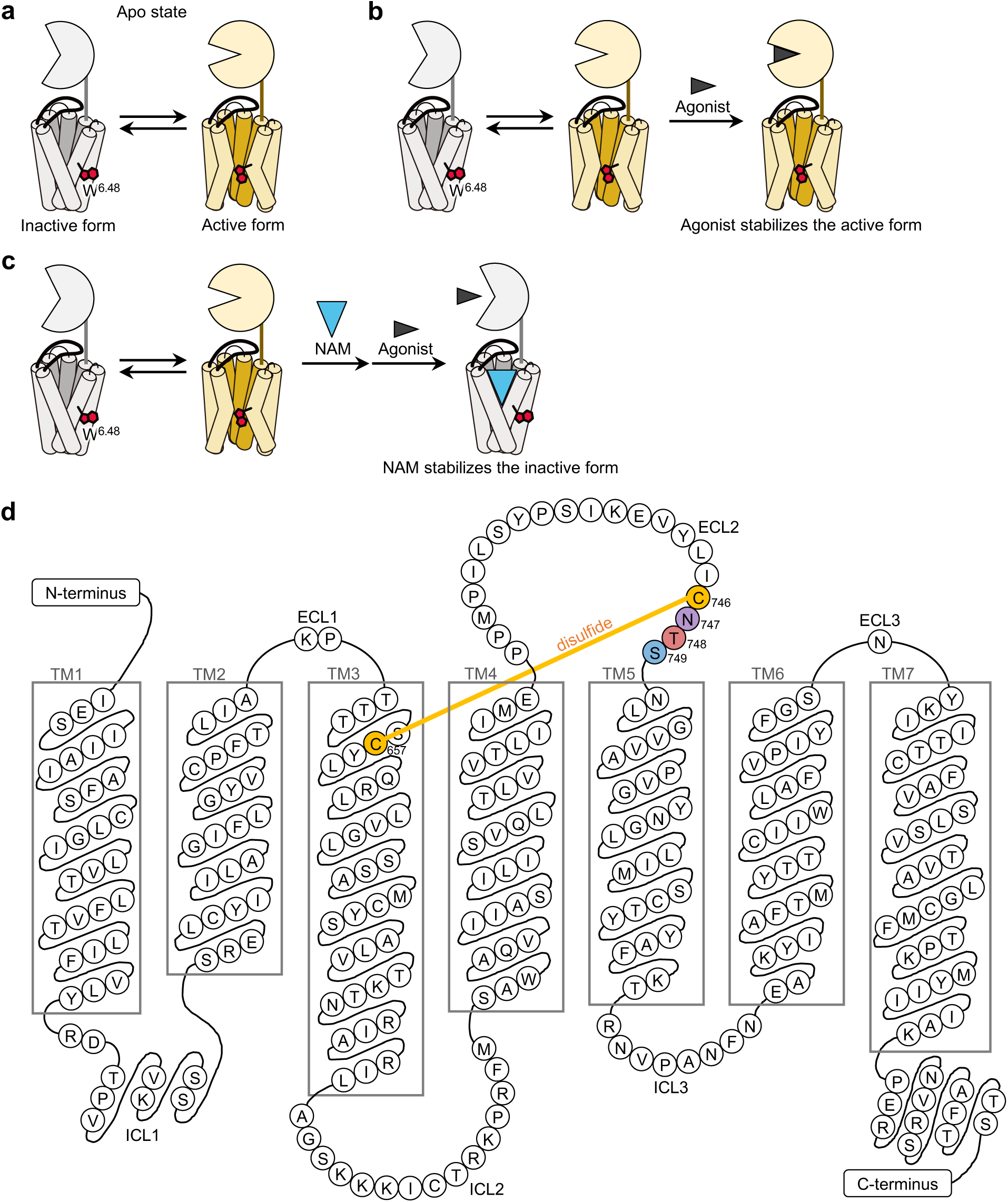
Dynamic equilibrium among conformational states of class C GPCRs. **a–c,** Schematic illustration of the apo (in **a**), agonist-bound (in **b**), and NAM-bound (in **c**) conformational states of mGlu1. **d,** Amino acid sequence of the 7-transmembrane domain (7TMD) of mouse mGlu1.

**Extended Data Fig. 2.**
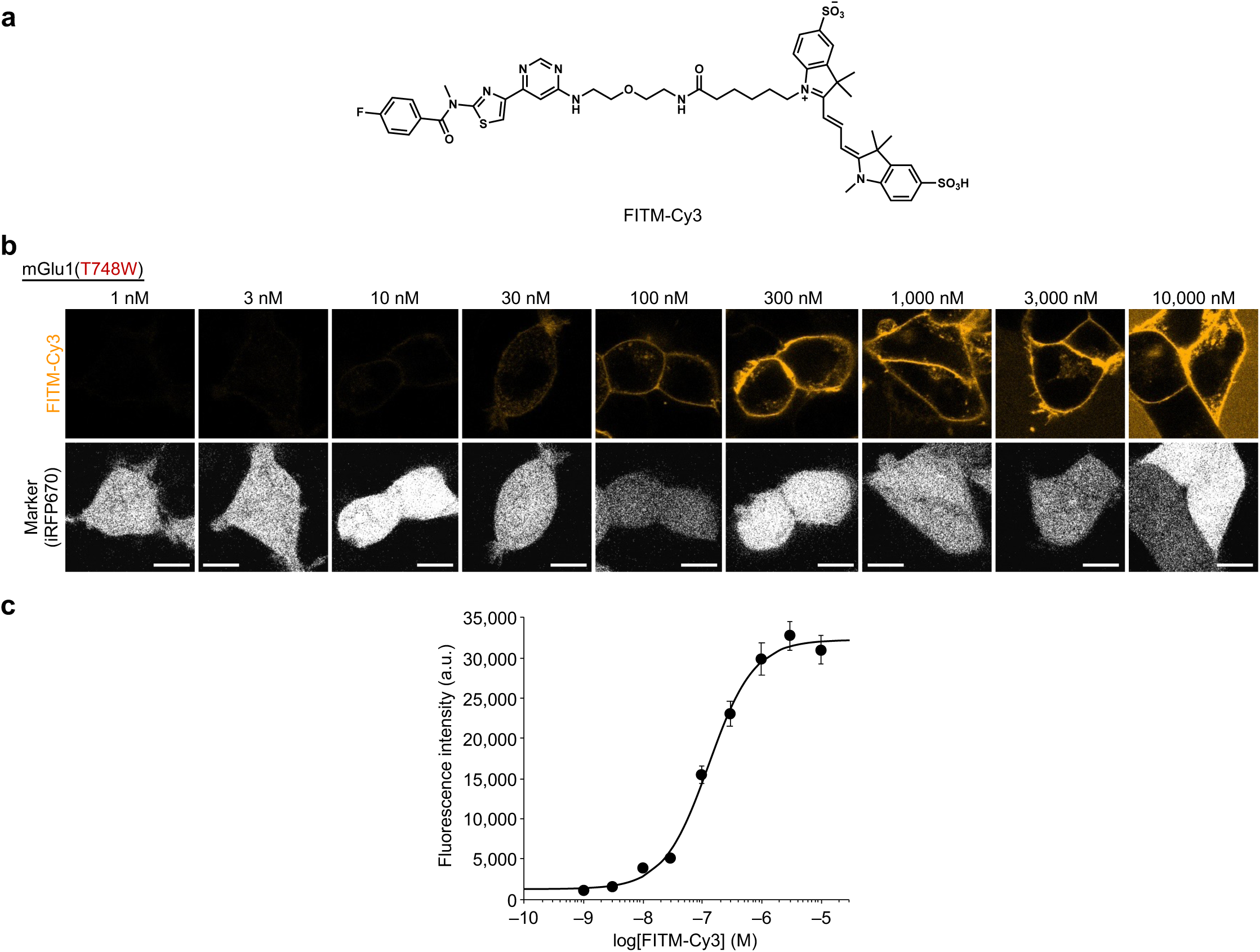
Quantification of FITM-Cy3 binding to mGlu1(T748W) by confocal microscopy. **a,** Chemical structure of FITM-Cy3. **b,** Live-cell confocal imaging of cell-surface mGlu1 visualized with increasing concentrations of FITM-Cy3 in HEK293 cells expressing mGlu1(T748W) (upper). iRFP670 was used as a transfection marker (lower). Scale bars, 10 µm. **c,** Concentration-dependent curves for fluorescence intensity of FITM-Cy3 in HEK293 cells expressing mGlu1(T748W) (*n* = 105 cells). The *K*_d_ of FITM-Cy3 for mGlu1(T748W) was determined to be 142 ± 29 nM. The *K*_d_ of FITM-Cy3 for WT mGlu1 was 6.8 ± 2.7 nM in a previous study^20^. Data are presented as mean ± SEM (*N* = 3).

**Extended Data Fig. 3.**
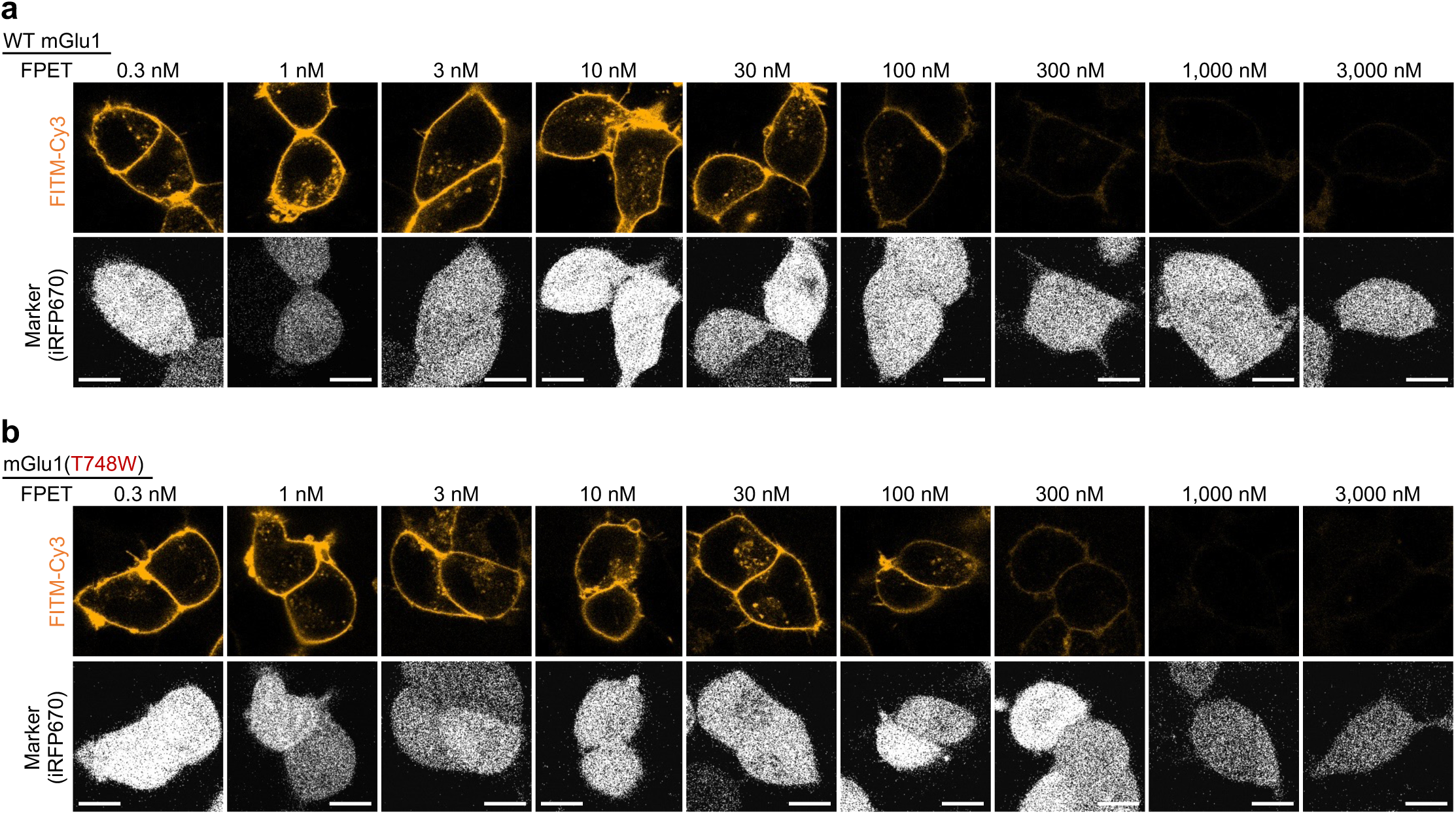
Competitive binding of FITM-Cy3 and FPET to WT mGlu1 and mGlu1(T748W). **a,b,** Live-cell confocal imaging of cell-surface mGlu1 visualized with FITM-Cy3 (30 nM for WT mGlu1 in **a**; 300 nM FITM-Cy3 for mGlu1(T748W) in **b**) with increasing concentrations of FPET in HEK293 cells expressing either WT mGlu1 or mGlu1(T748W) (upper). iRFP670 was used as a transfection marker (lower). Scale bars, 10 µm.

**Extended Data Fig. 4.**
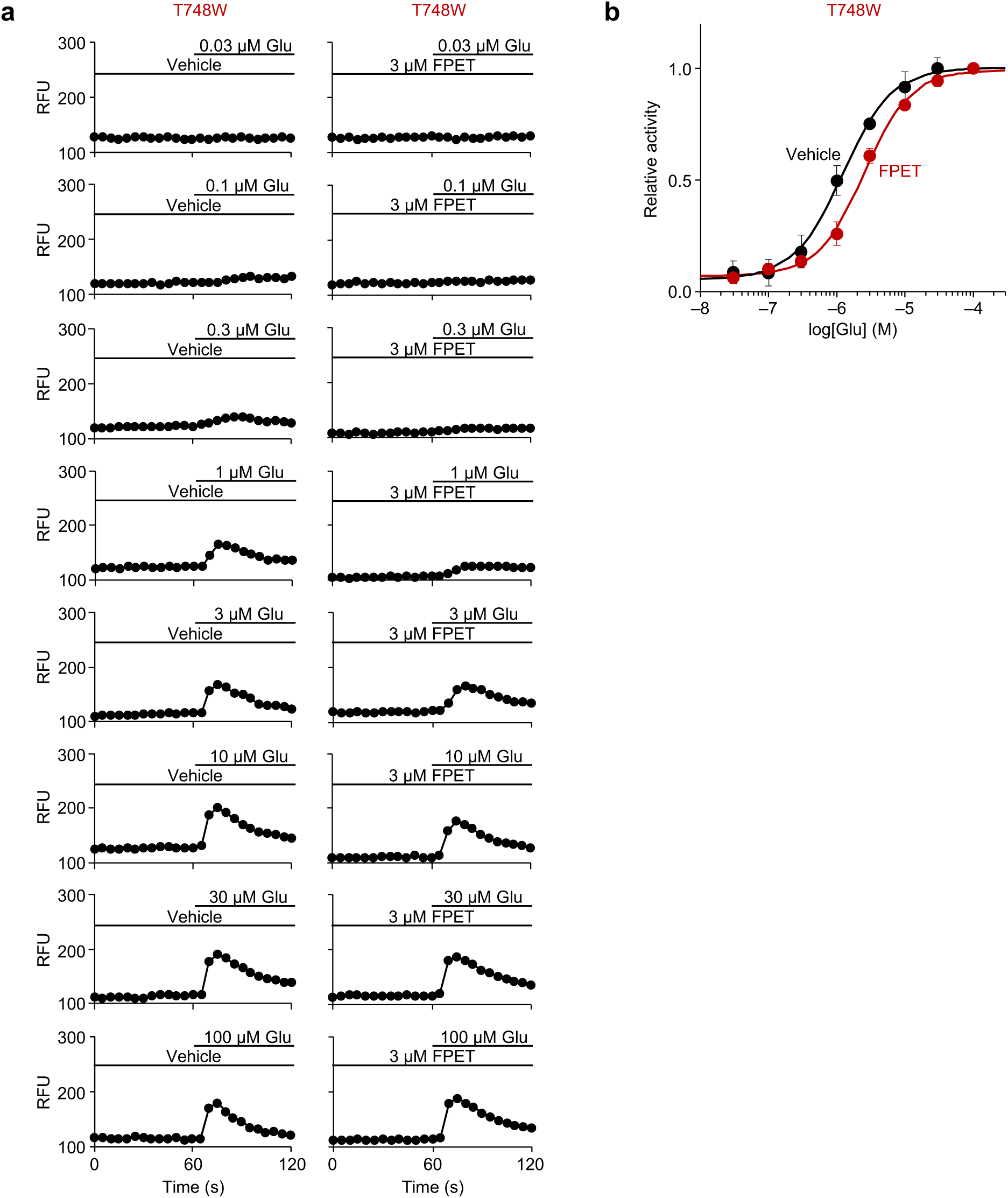
Glutamate-induced activation of mGlu1(T748W) in the presence or absence of 3 µM FPET. **a,** Representative Cal-520 fluorescence traces in HEK293 cells expressing mGlu1(T748W). Cells were pretreated with vehicle or 3 µM FPET for 3 min and subsequently stimulated with increasing concentrations of glutamate (Glu). **b,** The corresponding Glu concentration–response curves. Relative activity was calculated as the ratio of the obtained response intensity to that of mGlu1-expressing HEK293 cells induced by 100 µM Glu. The EC_50_ values for Glu-induced activation of mGlu1(T748W) were 1.2 ± 0.1 µM in the absence and 2.4 ± 0.4 µM in the presence of 3 µM FPET. Data are presented as mean ± SEM (*N* = 3).

**Extended Data Fig. 5.**
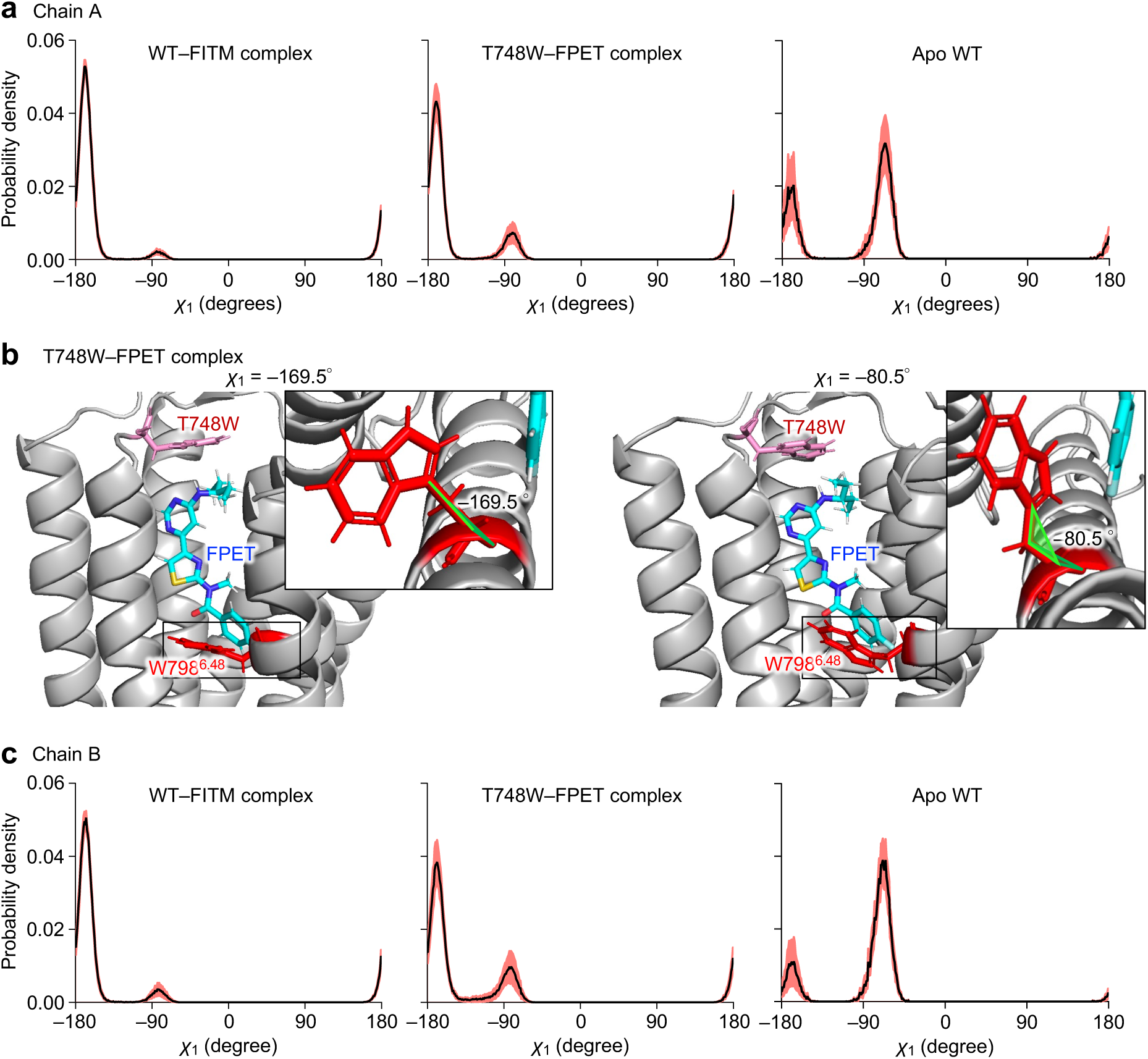
Comparison of χ_1_ dihedral angle distribution of W798^6^.^48^ in WT mGlu1–FITM complex, mGlu1(T748W)–FPET complex, and WT mGlu1 in the apo state. **a,** χ_1_ distribution of W798^6.48^ in the chain A of WT mGlu1–FITM complex (left), mGlu1(T748W)–FPET complex (middle), and WT mGlu1 in the apo state (right). **b,** The structure of mGlu1(T748W)–FPET complex at χ_1_ = –169.5°(left) and –80.5°(right). **c,** χ_1_ distribution of W798^6.48^ in the chain B of WT mGlu1–FITM complex (left), mGlu1(T748W)–FPET complex (middle), and WT mGlu1 in the apo state (right). The mean is shown as a black line, and ± SEM is shown in a rose color.

**Extended Data Fig. 6.**
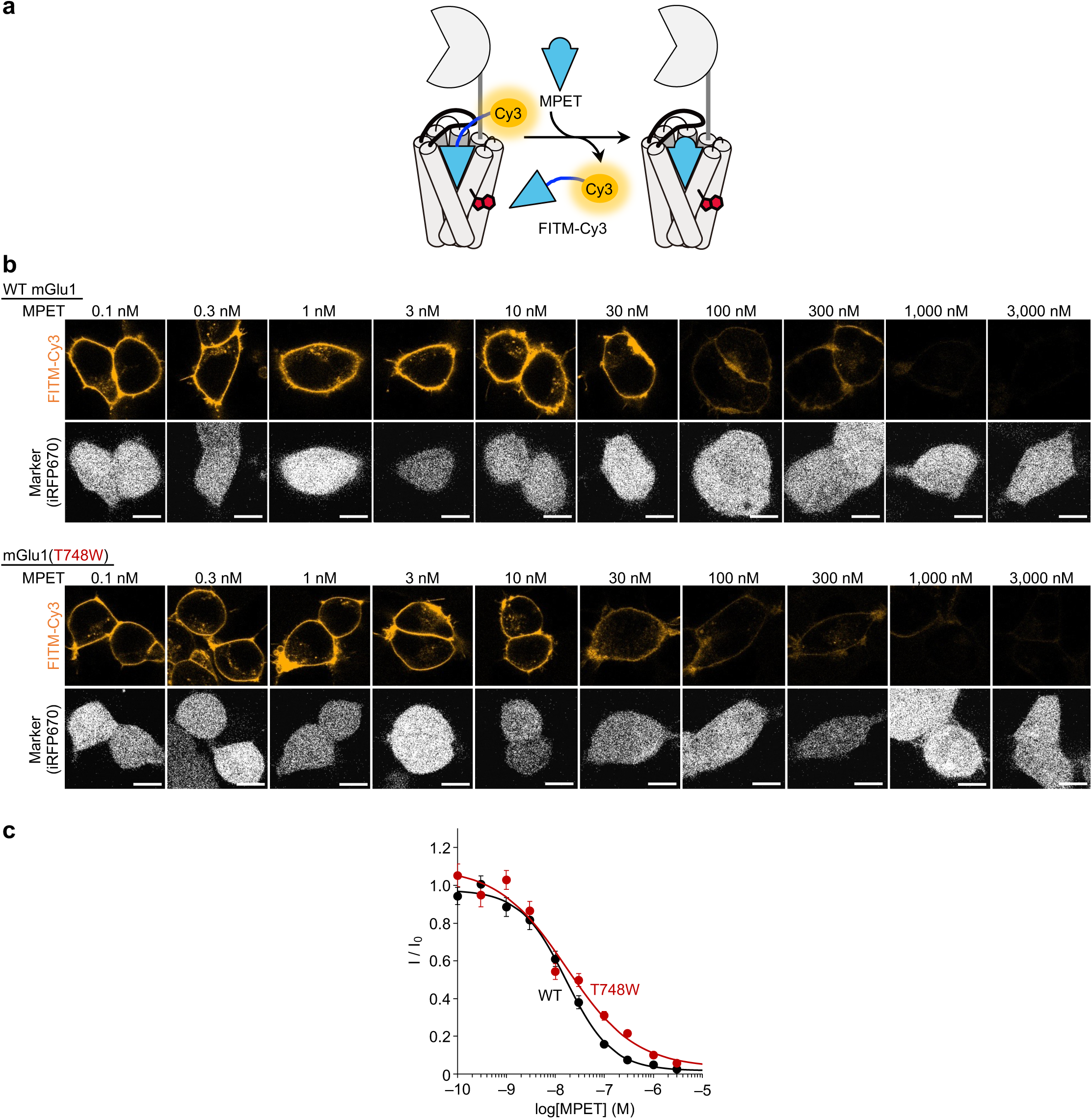
Competitive binding of FITM-Cy3 and MPET to WT mGlu1 and mGlu1(T748W). **a,** Schematic illustration of competitive binding of FITM-Cy3 and MPET to mGlu1. **b,** Live-cell confocal imaging of cell-surface mGlu1 visualized with FITM-Cy3 (30 nM for WT mGlu1 or 300 nM for mGlu1(T748W)) with increasing concentrations of MPET in HEK293 cells expressing WT mGlu1 or mGlu1(T748W) (upper). iRFP670 was used as a transfection marker (lower). Scale bars, 10 µm. **c,** Concentration-dependent displacement of FITM-Cy3 by MPET in HEK293 cells expressing WT mGlu1 or mGlu1(T748W) (*n* = 105 cells). Normalized fluorescence intensity (I/I_0_) was calculated relative to the fluorescence intensity of FITM-Cy3 obtained in the absence of MPET (I_0_). *K*_i_ values of MPET were 2.7 ± 0.6 nM for WT mGlu1 and 5.9 ± 1.8 nM for mGlu1(T748W). Data are presented as mean ± SEM (*N* = 3).

**Extended Data Fig. 7.**
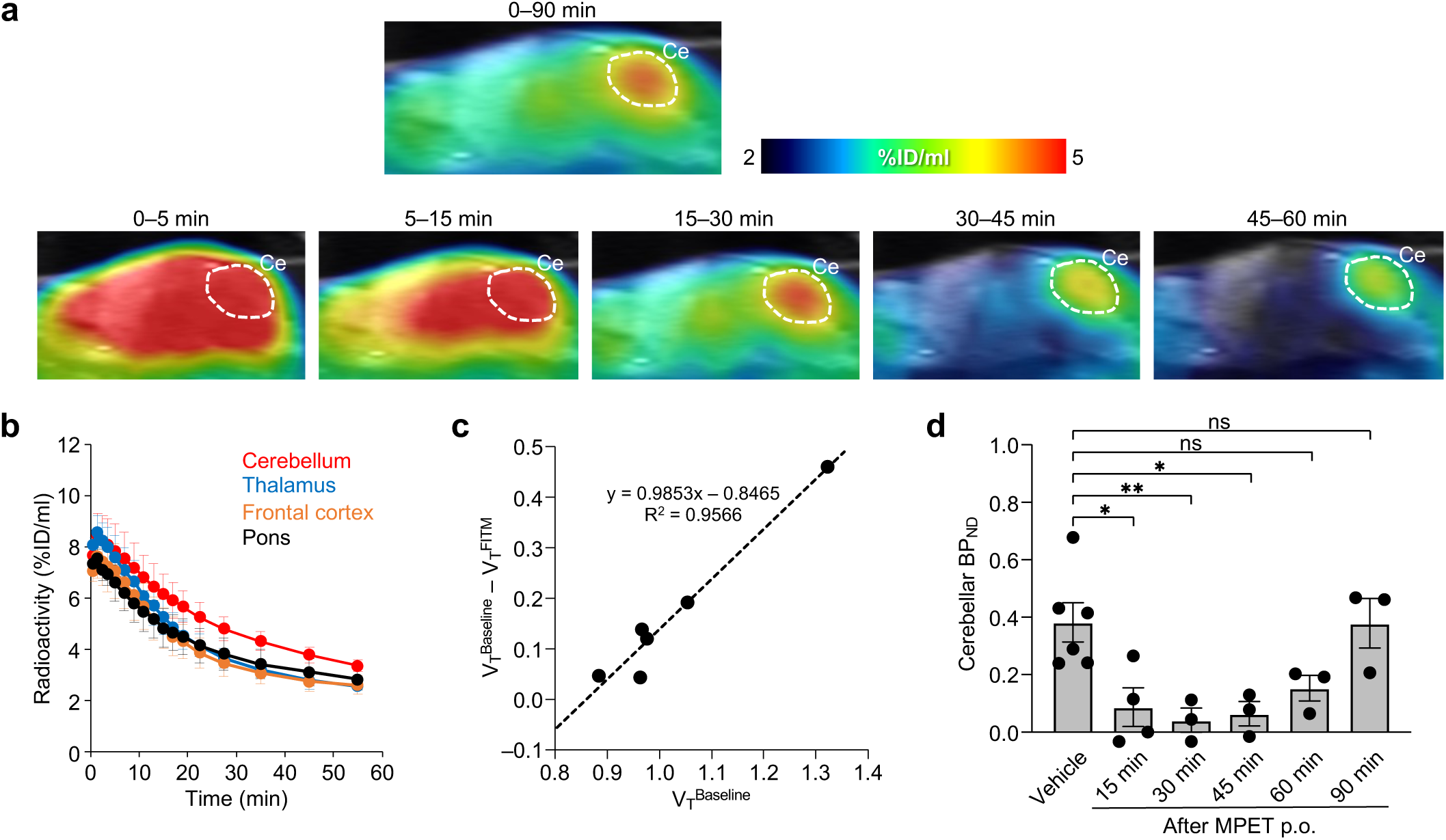
*In vivo* PET imaging of mouse brain using [^11^C]MPET. **a,** Upper, summed PET images, co-registered to an MRI template, acquired from 0 to 90 min after intravenous injection of [^11^C]MPET via the tail vein. Lower, time-dependent changes in radioactivity uptake following intravenous injection of [^11^C]MPET. Ce, cerebellum. **b,** Time-activity curves (TACs) of [^11^C]MPET in the cerebellum, thalamus, frontal cortex, and pons of mice orally pretreated with vehicle 30 min before injection. Radioactivity is expressed as percentage of injected dose per milliliter (%ID/ml). **c,** Lassen plots generated using regional distribution volume (V_T_) values under baseline conditions and full mGlu1 blockade (pretreatment with FITM). The slope represents FITM occupancy at the mGlu1 binding site, and the y-axis intercept indicates the non-displaceable distribution volume (V_ND_) values of [^11^C]MPET in mouse brain. **d,** Cerebellar BP_ND_ (V_T_/V_ND_ − 1) values of [^11^C]MPET in mice orally pre-administered with MPET (6 mg/kg) 15, 30, 45, 60, or 90 min before injection. Data are presented as mean ± SEM (*n* = 3–6 mice per group). Statistical analyses were performed using Dunnett’s test compared with the vehicle group. \**P* < 0.05, \*\**P* < 0.01, ns, not significant.

**Extended Data Fig. 8.**
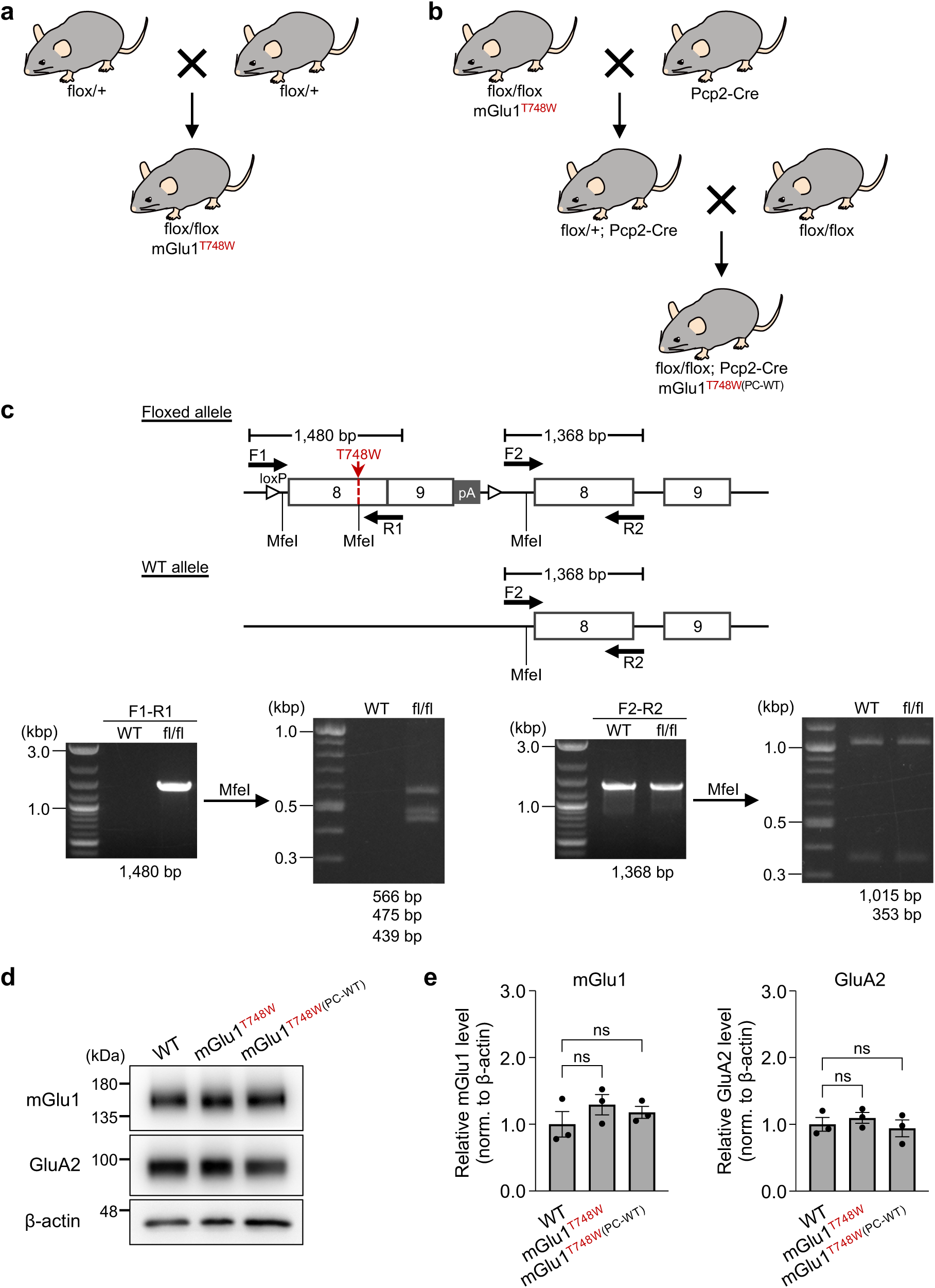
Generation and validation of *GRM1* conditional knock-in mice harboring the T748W mutation. **a,** Strategy for generating floxed (fl/fl) knock-in mice expressing mGlu1(T748W) (mGlu1^T748W^ mice). **b,** Generation of mGlu1^T748W^ mice expressing Cre recombinase under the control of the Purkinje cell-specific Pcp2 promoter, resulting in Purkinje cell-specific reversion to WT mGlu1 (mGlu1^T748W(PC-WT)^ mice). **c,** Genotyping strategy for mGlu1^T748W^ mice. The T748W mutation introduces a MfeI restriction site into the floxed allele. Genotyping was performed by PCR using F1/R1 or F2/R2 primer pairs followed by MfeI digestion. **d,** Western blot analysis of mGlu1, GluA2, and β-actin in cerebellar lysates from WT, mGlu1^T748W^, and mGlu1^T748W(PC-WT)^ mice. **e,** Quantification of the mGlu1 and GluA2 expression in the cerebellum. Data are presented as mean ± SEM (*n* = 3 mice per group). ns, not significant. Statistical analysis were performed using Dunnett’s test compared with the WT control.

**Extended Data Fig. 9.**
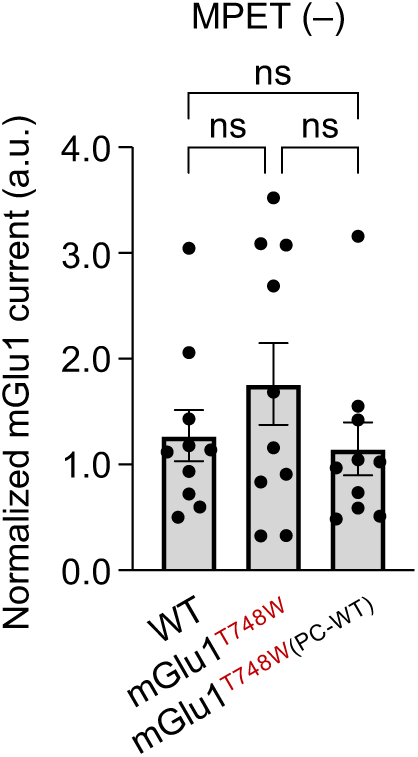
mGlu1-mediated slow currents recorded from cerebellar Purkinje cells in WT, mGlu1^T748W^, and mGlu1^T748W(PC-WT)^ mice. mGlu1-mediated slow currents were normalized to the peak amplitude of AMPAR-mediated fast currents. Data are presented as mean ± SEM (*n* = 10 cells per mouse). ns, not significant. Statistical analyses were performed using the Kruskal–Wallis test.

**Extended Data Fig. 10.**
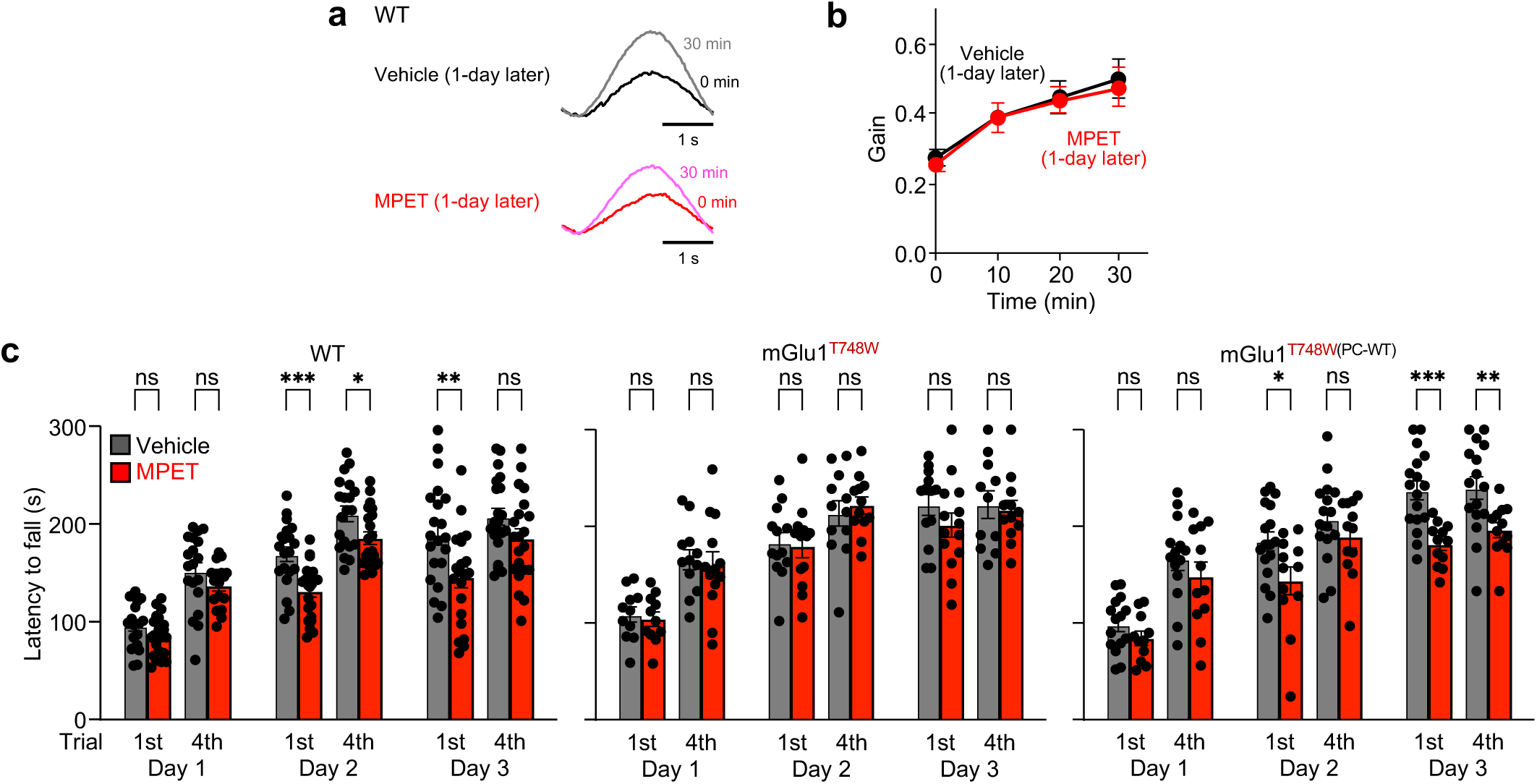
Reversibility of NARCH-mediated mGlu1 modulation and analysis of accelerating rotarod performance. **a,** Representative hOKR waveforms recorded before training (0 min) and 30 min after the onset of training. **b,** Averaged hOKR adaptation in WT mice following oral administration of vehicle or MPET (6 mg/kg) 1 day before hOKR testing. Data are presented as the mean ± SEM (*n* = 10 mice per group). **c**, Latency to fall in the first and last trials on day 1, 2, and 3 in the accelerating rotarod test. Data are presented as mean ± SEM (WT: *n* = 19–21 vehicle, 17–22 MPET; mGlu1^T748W^, *n* = 11–13 vehicle, 11–14 MPET; mGlu1^T748W(PC-WT)^: *n* = 14–17 vehicle, 11–12 MPET). \*\*\**p* < 0.001; \*\**p* < 0.01; \**p* < 0.05; ns, not significant. Statistical analyses were performed using the unpaired *t*-test.

## Supplementary Information

**Supplementary Table 1.**
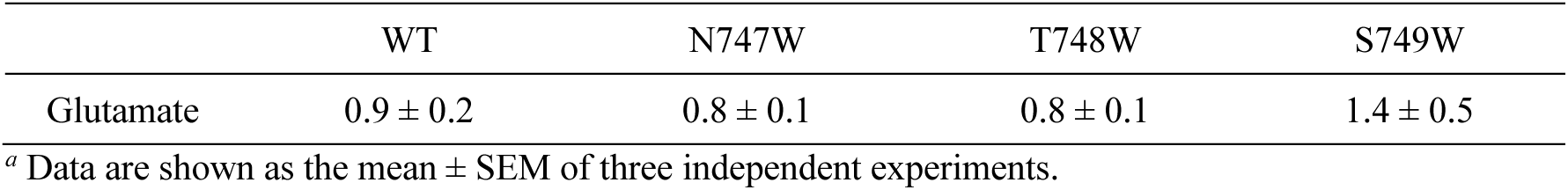
EC_50_ values (µM) of glutamate for mGlu1 (WT, N747W, T748W or S749W) *^a^*.

|  | WT | N747W | T748W | S749W |
| --- | --- | --- | --- | --- |
| Glutamate | 0.9 ± 0.2 | 0.8 ± 0.1 | 0.8 ± 0.1 | 1.4 ± 0.5 |

<sup>a</sup> Data are shown as the mean ± SEM of three independent experiments.

**Supplementary Table 2.** IC_50_ values (nM) of FITM for mGlu1 (WT, N747W, T748W or S749W) *^a,b^*.

|  | WT | N747W | T748W | S749W |
| --- | --- | --- | --- | --- |
| FITM | 3.3 ± 0.3 | 2.8 ± 0.8 | 17 ± 3.3 | 7.7 ± 2.3 |

<sup>a</sup> Data are shown as the mean ± SEM of three independent experiments.
<sup>b</sup> mGlu1 expressed in HEK293 cells was stimulated by 10 μM glutamate.

**Supplementary Fig. 1.**
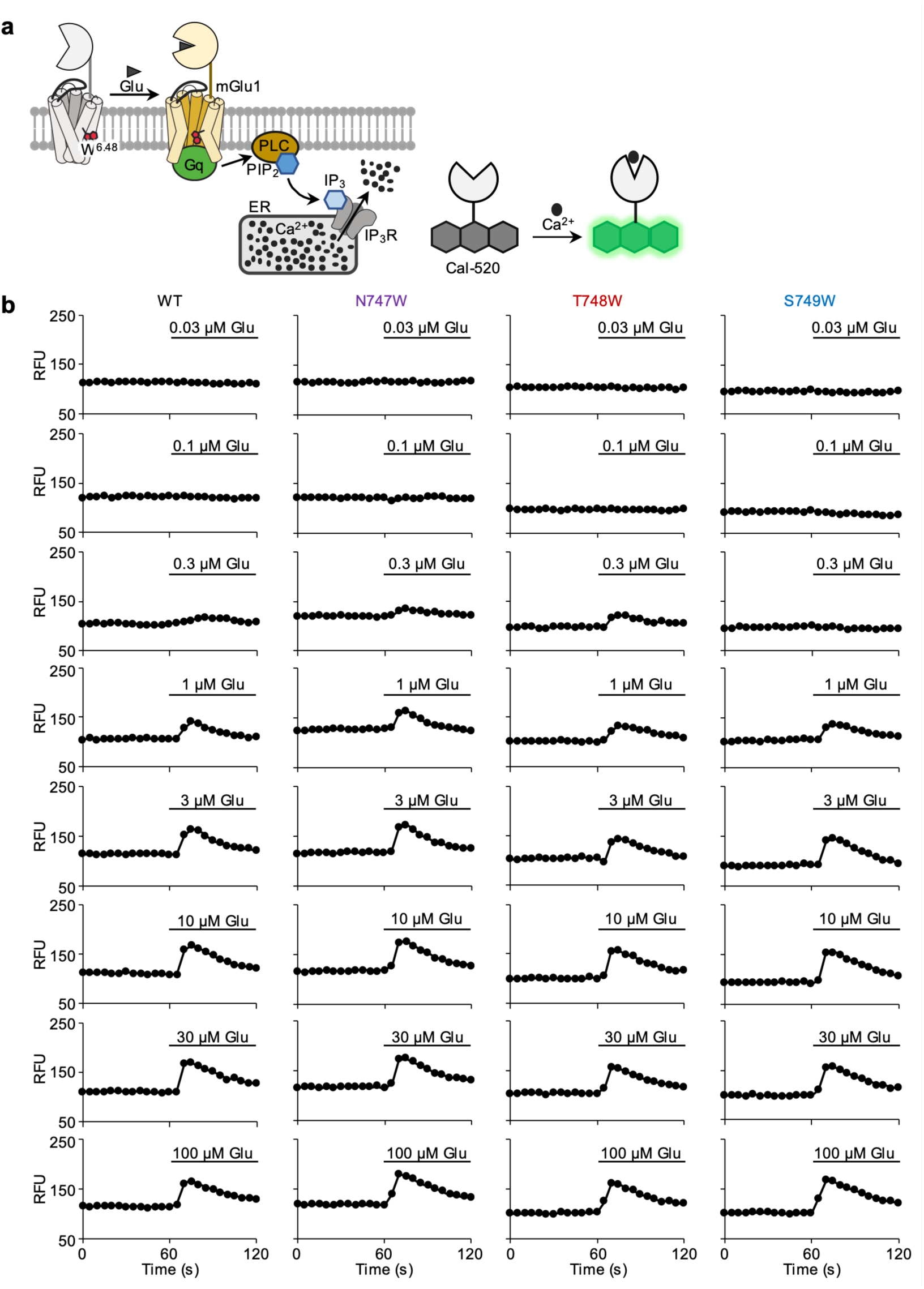
Activation of mGlu1 mutants by glutamate. **a,** Schematic illustration of glutamate (Glu)-induced mGlu1 signaling. The resulting increase in intracellular Ca^2+^ was detected using the fluorescent Ca^2+^ indicator Cal-520. **b,** Cal-520 fluorescence signals in HEK293 cells expressing mGlu1 (WT, N747W, T748W, or S749W) were recorded after stimulation with different concentrations of Glu. Data are presented as relative fluorescence units (RFU).

**Supplementary Fig. 2.**
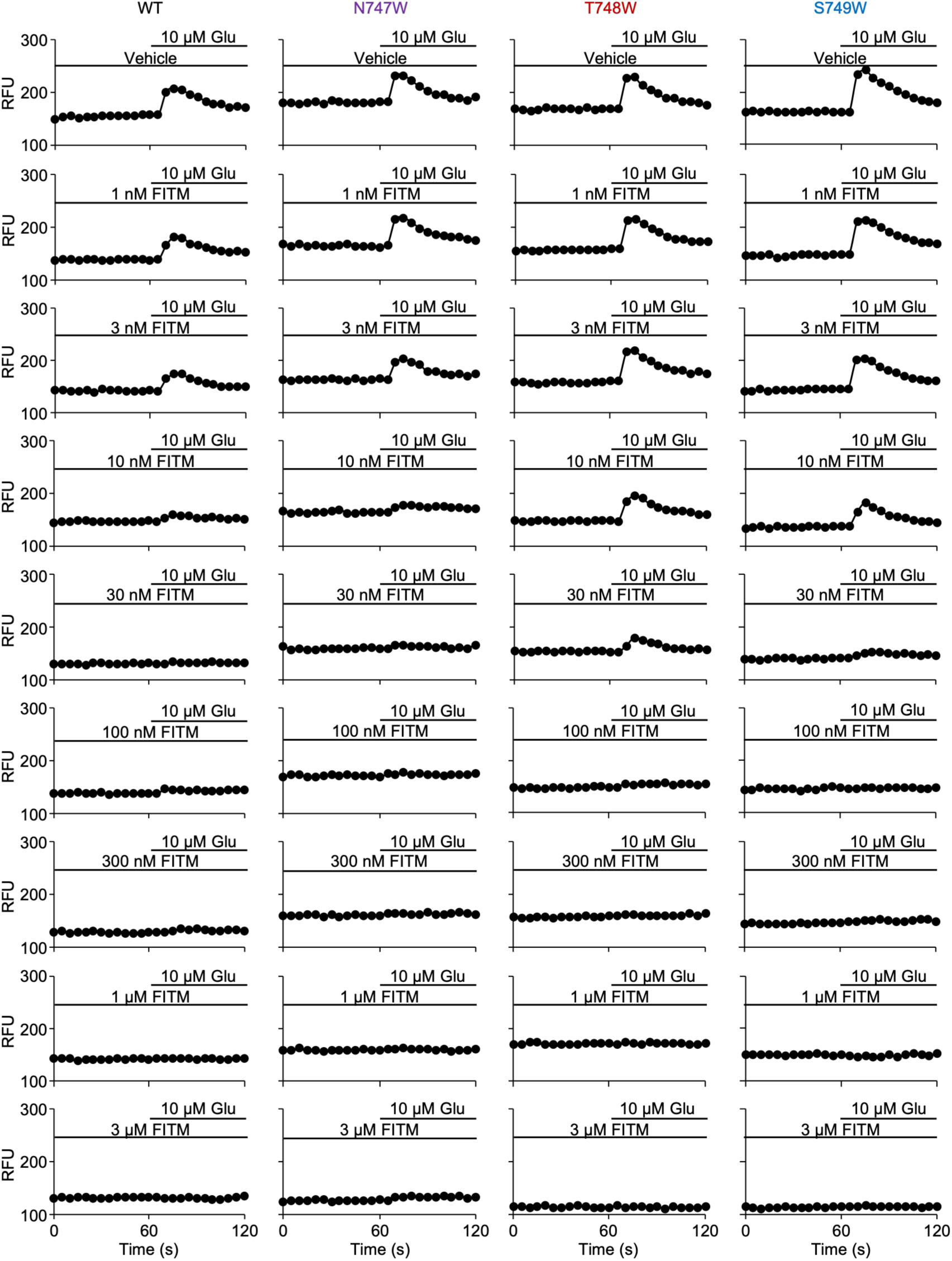
Inhibition of mGlu1 mutants by FITM. Cal-520 fluorescence signals were measured in HEK293 cells expressing mGlu1 (WT, N747W, T748W, or S749W). Cells were pretreated with FITM at different concentrations for 3 min and then stimulated with 10 µM Glu. Data are presented as RFU.

**Supplementary Fig. 3.**
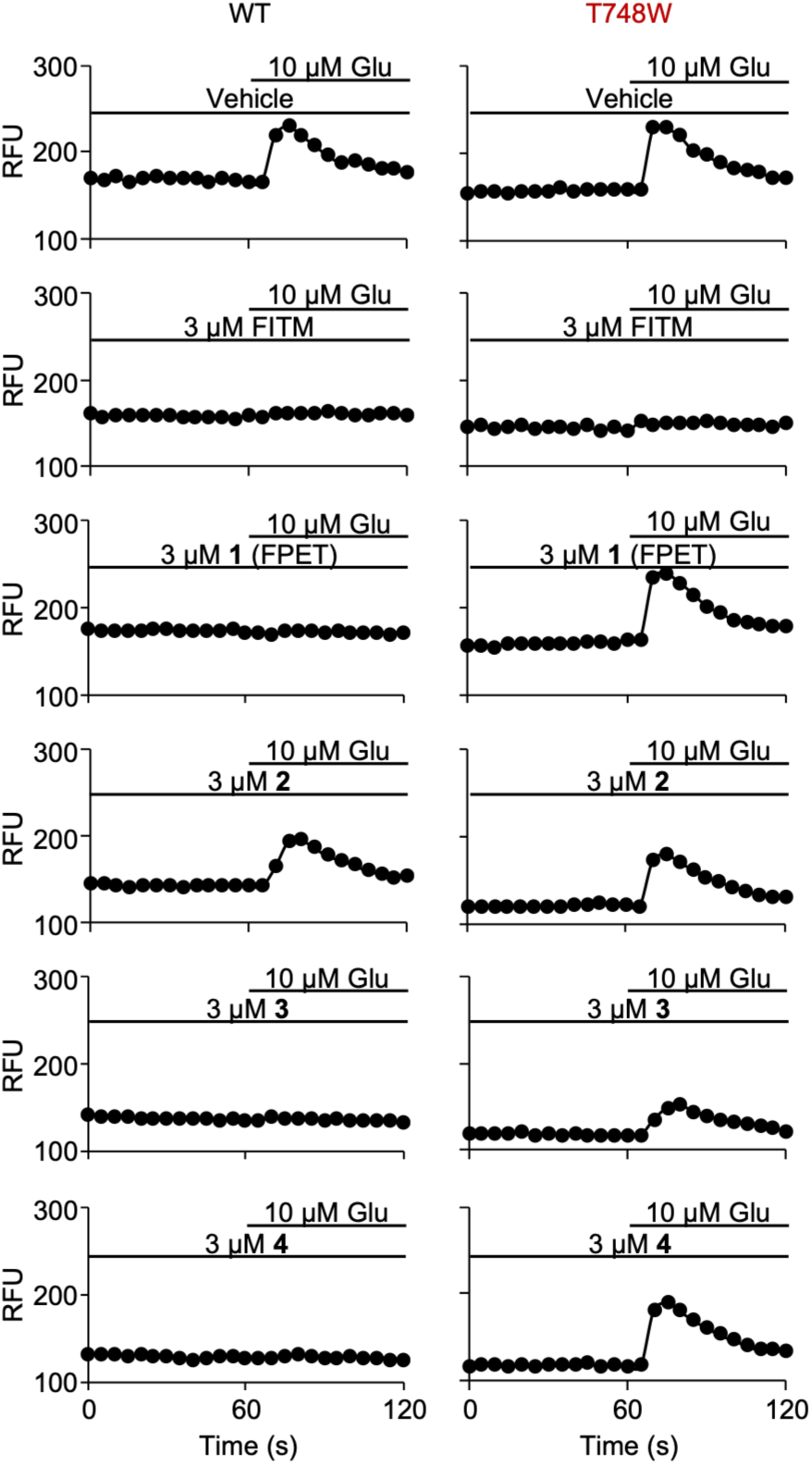
Inhibition of WT mGlu1 and mGlu1(T748W) by FITM derivatives 1–4. Cal-520 fluorescence signals were measured in HEK293 cells expressing mGlu1 (WT or T748W). Cells were pretreated with 3 µM FITM derivatives **1**–**4** for 3 min and then stimulated with 10 µM Glu. Data are presented as RFU.

**Supplementary Fig. 4.**
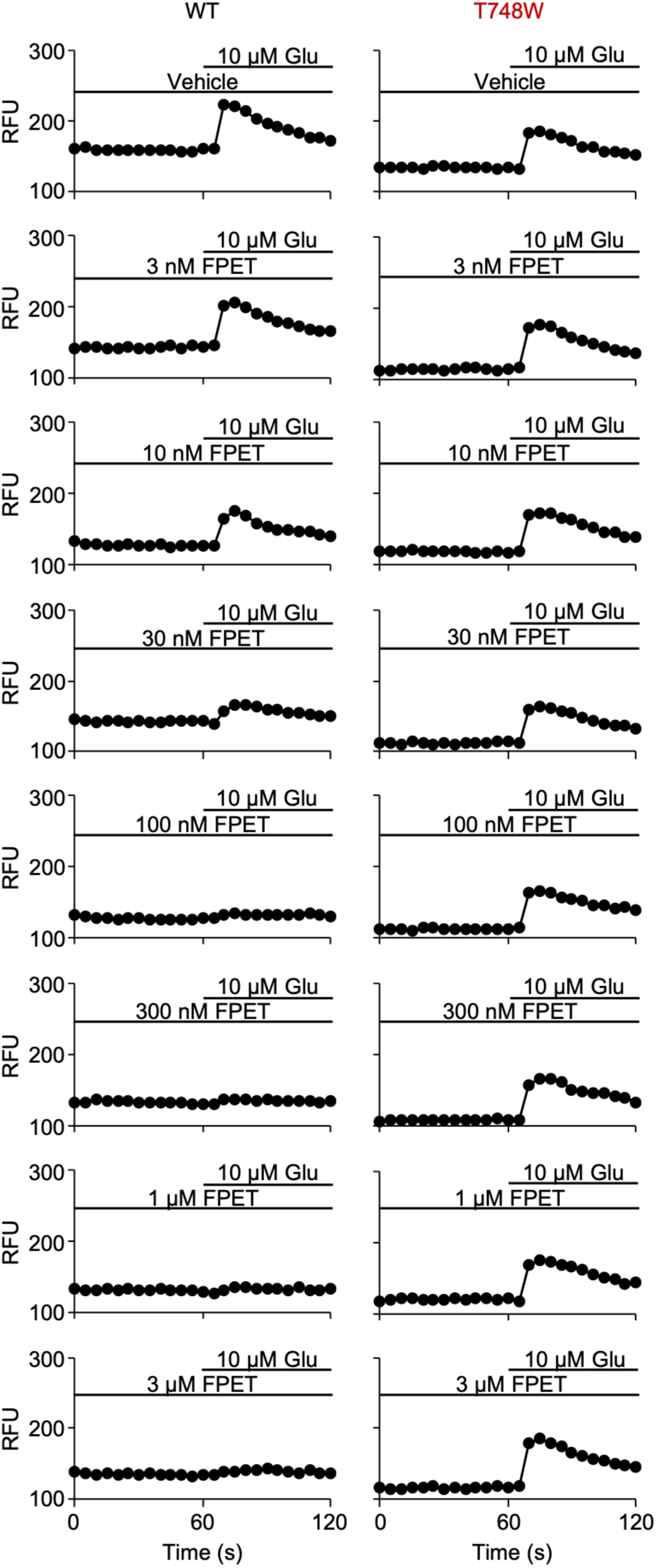
Inhibition of WT mGlu1 and mGlu1(T748W) by FPET. Cal-520 fluorescence signals were measured in HEK293 cells expressing mGlu1 (WT or T748W). Cells were pretreated with different concentrations of FPET for 3 min and then stimulated with 10 µM Glu. Data are presented as RFU.

**Supplementary Fig. 5.**
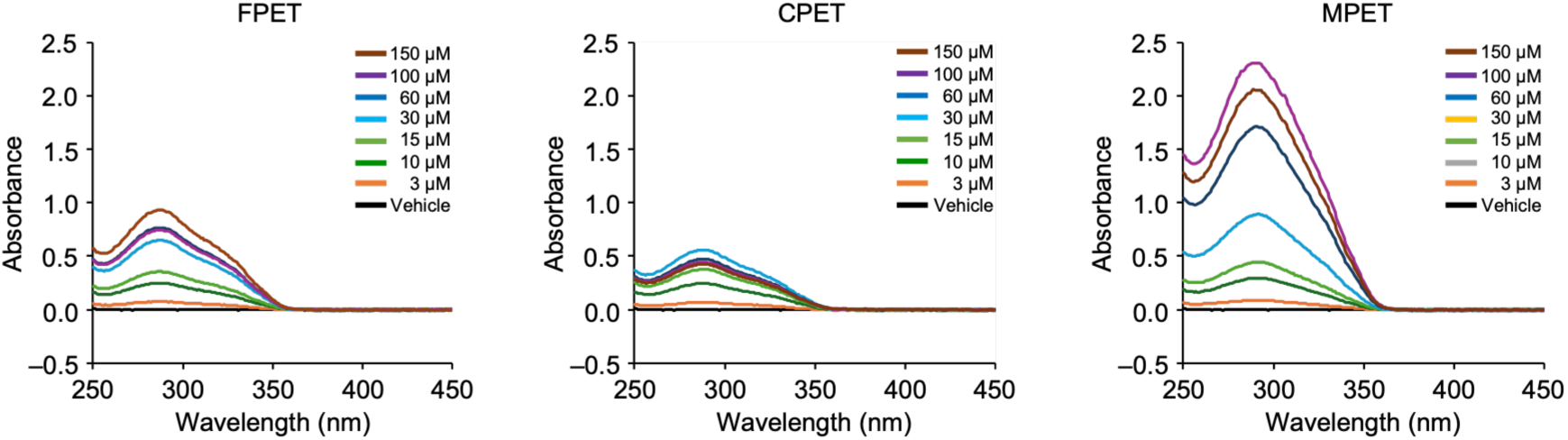
UV–Vis spectra of FPET, CPET, and MPET. Solutions of FPET, CPET, and MPET in DMSO at different concentrations were added to 100 mM GTA buffer (pH 3.5) to mimic the acidic environment of the mouse stomach, and incubated for 1 h at 25 ℃. After incubation, UV–Vis spectra of the supernatants were measured.

**Supplementary Fig. 6.**
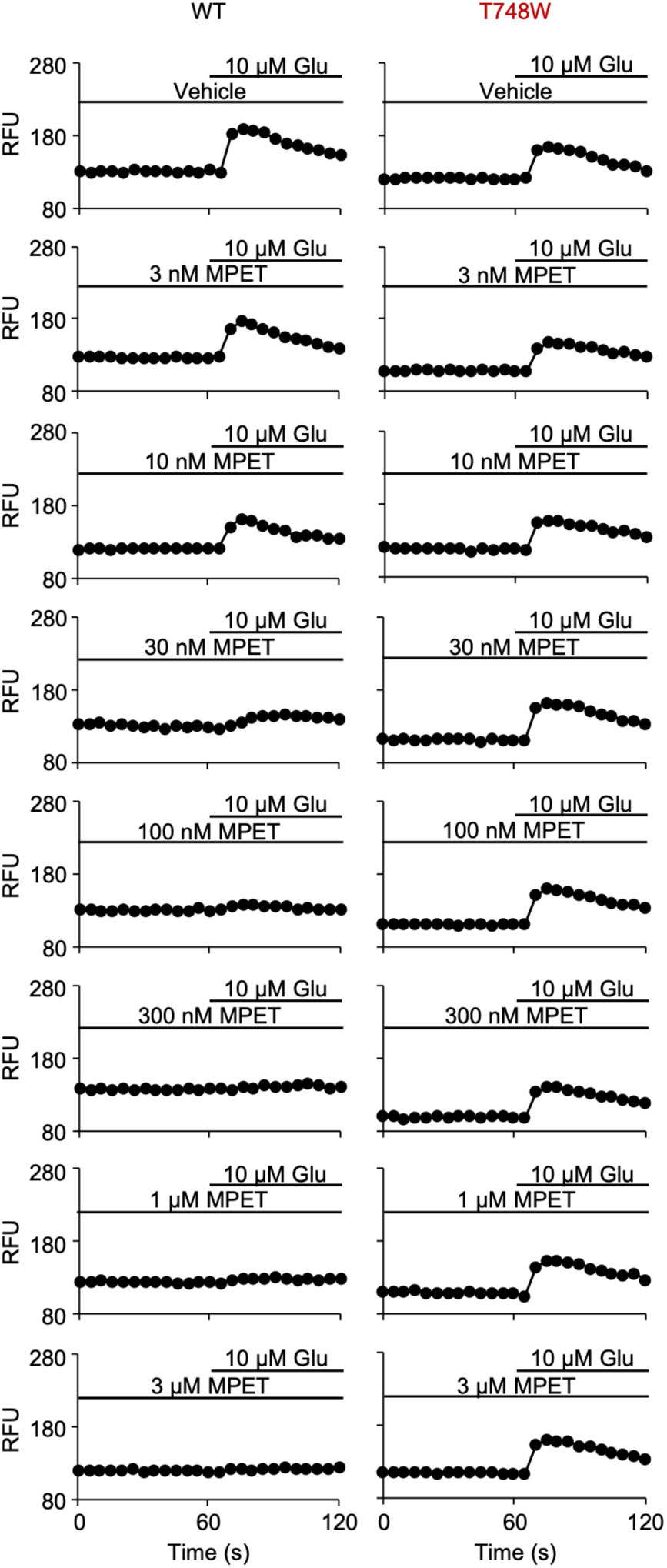
Inhibition of WT mGlu1 and mGlu1(T748W) by MPET. Cal-520 fluorescence signals were measured in HEK293 cells expressing mGlu1 (WT or T748W). Cells were pretreated with different concentrations of MPET for 3 min and then stimulated with 10 µM Glu. Data are presented as RFU.

## Supplementary Notes (Chemical Synthesis)

### Materials and instruments for organic synthesis

All chemical reagents and solvents were purchased from commercial suppliers (Tokyo Chemical Industry (TCI), Fujifilm-Wako Pure Chemical Corporation, Kanto Chemical Co., Inc., Sigma-Aldrich, and Lumiprobe) and used without further purification. Thin-layer chromatography (TLC) was performed on silica gel 60 F254 precoated glass plates (Merck Millipore) and visualized by fluorescence quenching and ninhydrin staining. Preparative thin layer chromatography (PLC) was carried out using PLC silica gel 60 F_254_ (1 mm, 20 × 20 cm, Merck Millipore). Chromatographic purification was carried out using Wakosil C-200 (spherical, 64–210 µm, Fujifilm-Wako) and silica gel 60 N (spherical, neutral, 40–50 µm, Kanto Chemical). ^1^H NMR (300 MHz or 500 MHz) spectra were recorded in deuterated solvents on a Bruker AVANCE III HD 300 MHz or 500 MHz spectrometer. Chemical shifts (*δ*, ppm) were referenced to tetramethylsilane (0 ppm) or residual solvent signals including CD_2_HOD (3.30 ppm, MeOH-*d*_4_), CD_2_HSOCD_3_ (2.50 ppm, DMSO-*d*_6_), and CHCl_2_CDCl_2_ (6.00 ppm, 1,1,2,2-tetrachloroethane-*d*_2_). Multiplicities are abbreviated as follows: s = singlet, brs = broad singlet, d = doublet, brd = broad doublet, t = triplet, dd = doublet of doublets, m = multiplet, brm = broad multiplet. Matrix-assisted laser desorption/ionization time-of-flight mass spectrometry (MALDI-TOF MS) spectra were recorded on an autoflex maX instrument (Bruker Daltonics) using dithranol (DIT) as the matrix. High-resolution mass spectra (HRMS) were acquired on a compact mass spectrometer (Bruker Daltonics) equipped with electrospray ionization (ESI).

### Synthesis of FITM derivatives

Compound **7** and FITM were synthesized according to previously reported procedures with modifications (ref. S1). Compounds **8** and **9** were synthesized according to modified literature procedures (refs. S2 and S3), respectively.

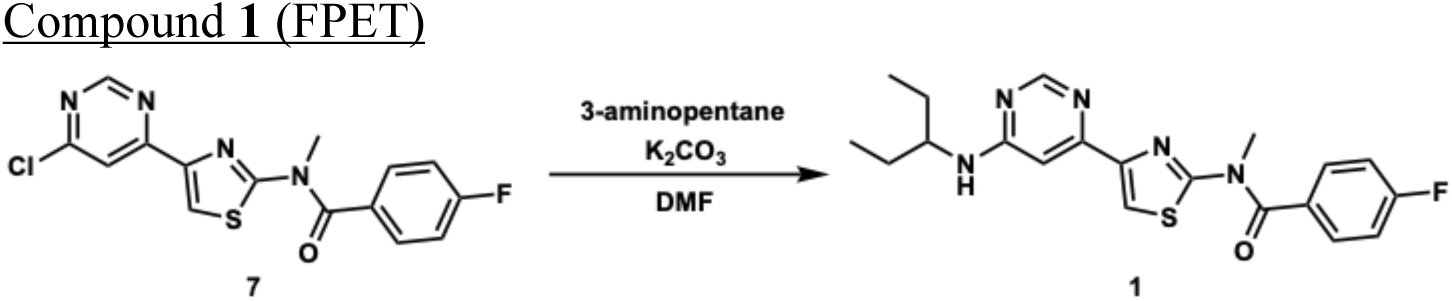

3-Aminopentane (40 µl, 0.344 mmol, 2.4 equiv.) was added to a mixture of compound **7** (50 mg, 0.143 mmol, 1 equiv.) and K_2_CO_3_ (27 mg, 0.195 mmol, 1.4 equiv.) in dry DMF (5 ml). The reaction mixture was stirred under N_2_ at 90 °C overnight. The mixture was concentrated *in vacuo* to remove the solvent, and the residue was diluted with EtOAc. The organic layer was washed twice with saturated aqueous NH_4_Cl and brine, dried over MgSO_4_, and concentrated *in vacuo*. The residue was purified by column chromatography (silica gel, CHCl_3_ to CHCl_3_/EtOAc = 50:1 to 25:1) followed by preparative thin-layer chromatography (silica gel, CHCl_3_/EtOAc = 1:1) to afford compound **1** (9 mg, 0.023 mmol, 16%) as a white solid. ^1^H NMR (500 MHz, 1,1,2,2-tetrachloroethane-*d*_2_, 353 K): *δ* 8.56 (d, *J* = 0.9 Hz, 1H), 7.97 (s, 1H), 7.63 (dd, *J* = 5.2, 8.8 Hz, 2H), 7.23 (t, *J* = 8.7 Hz, 2H), 7.09 (d, *J* = 0.9 Hz, 1H), 4.85 (brd, *J* = 8.3 Hz, 1H, NH), 3.81 (brm, 1H), 3.77 (s, 3H), 1.75–1.53 (m, 4H), 1.00 (t, *J* = 7.4 Hz, 6H). HRMS (ESI^+^) *m*/*z* Calcd. for C_20_H_23_FN_5_OS^+^ [M+H]^+^ 400.1602; Found 400.1607.

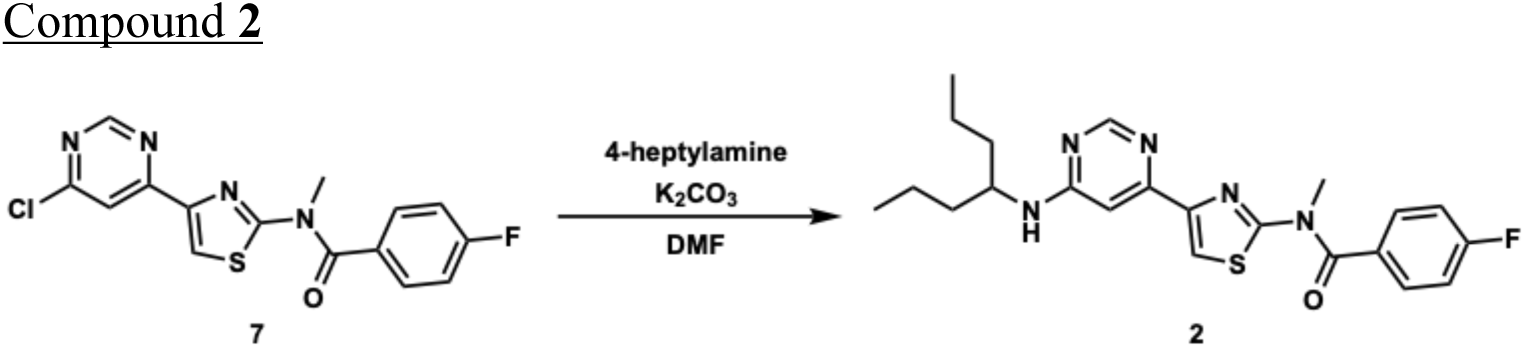

4-heptylamine (50 µl, 0.330 mmol, 2.3 equiv.) was added to a mixture of compound **7** (50 mg, 0.143 mmol, 1 equiv.) and K_2_CO_3_ (28 mg, 0.203 mmol, 1.4 equiv.) in dry DMF (5 ml). The reaction mixture was stirred under N_2_ at 95 °C for 7 h. The mixture was concentrated *in vacuo* to remove the solvent, and the residue was diluted with EtOAc. The organic layer was washed twice with saturated aqueous NH_4_Cl and brine, dried over MgSO_4_, and concentrated *in vacuo*. The residue was purified by column chromatography (silica gel, CHCl_3_ to CHCl_3_/EtOAc = 50:1) to afford compound **2** (16 mg, 0.037 mmol, 26%) as a white solid. ^1^H NMR (500 MHz, 1,1,2,2-tetrachloroethane-*d*_2_, 353 K): *δ* 8.55 (d, *J* = 0.8 Hz, 1H), 7.97 (s, 1H), 7.63 (dd, *J* = 5.2, 8.8 Hz, 2H), 7.23 (t, *J* = 8.7 Hz, 2H), 7.09 (d, *J* = 0.8 Hz, 1H), 4.76 (brd, *J* = 8.4 Hz, 1H, NH), 3.95 (brm, 1H), 3.77 (s, 3H), 1.67–1.40 (m, 8H), 1.00 (t, *J* = 7.2 Hz, 6H). HRMS (ESI^+^) *m*/*z* Calcd. for C_22_H_27_FN_5_OS^+^ [M+H]^+^ 428.1915; Found 428.1922.

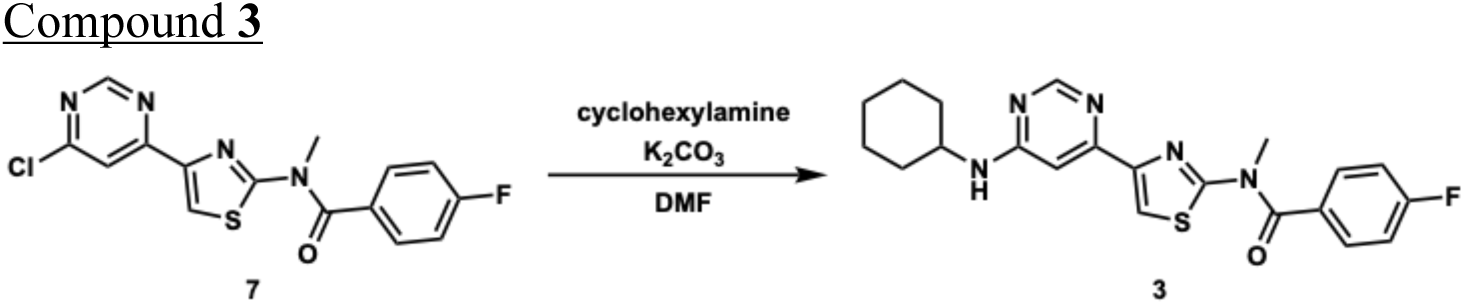

Cyclohexylamine (40 µl, 0.353 mmol, 2.5 equiv.) was added to a mixture of compound **7** (50 mg, 0.143 mmol, 1 equiv.) and K_2_CO_3_ (27 mg, 0.195 mmol, 1.4 equiv.) in dry DMF (5 ml). The reaction mixture was stirred under N_2_ at 90 °C for 9 h. The mixture was concentrated *in vacuo* to remove the solvent, and the residue was diluted with EtOAc. The organic layer was washed twice with saturated aqueous NH_4_Cl and brine, dried over MgSO_4_, and concentrated *in vacuo*. The residue was purified by column chromatography (silica gel, CHCl_3_ to CHCl_3_/EtOAc = 49:1 to 24:1) to afford compound **3** (32 mg, 0.078 mmol, 54%) as a white solid. ^1^H NMR (300 MHz, CDCl_3_): *δ* 8.56 (d, *J* = 1.1 Hz, 1H), 7.92 (s, 1H), 7.61 (dd, *J* = 5.2, 8.9 Hz, 2H), 7.20 (t, *J* = 8.9 Hz, 2H), 7.04 (d, *J* = 1.1 Hz, 1H), 4.96 (brs, 1H, NH), 3.75 (s, 3H), 2.09–2.04 (m, 1H), 1.82–1.76 (m, 4H), 1.51–1.38 (m, 2H), 1.32–1.20 (m, 4H). HRMS (ESI^+^) *m*/*z* Calcd. for C_21_H_23_FN_5_OS^+^ [M+H]^+^ 412.1602; Found 412.1606.

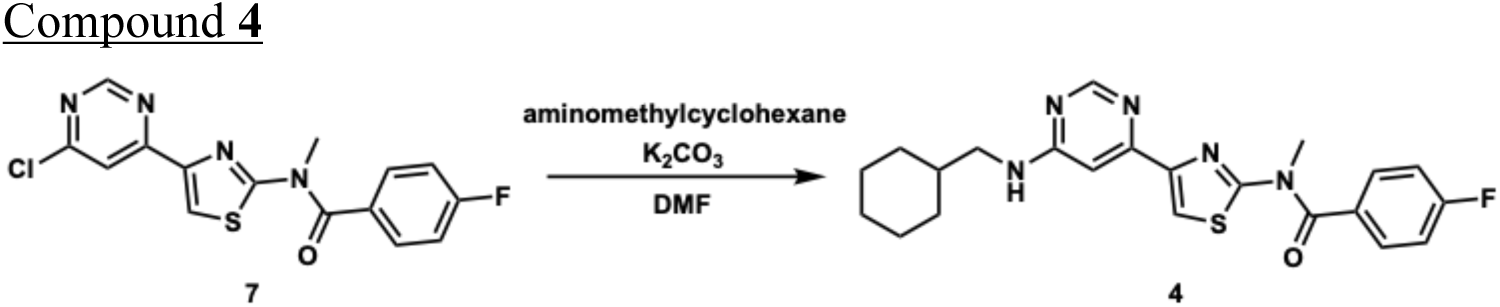

<u>A</u>minomethylcyclohexane (40 µl, 0.307 mmol, 2.2 equiv.) was added to a mixture of compound **7** (48 mg, 0.138 mmol, 1 equiv.) and K_2_CO_3_ (25 mg, 0.181 mmol, 1.3 equiv.) in dry DMF (4.8 ml). The reaction mixture was stirred under N_2_ at 95 °C for 6.5 h. The mixture was concentrated *in vacuo* to remove the solvent, and the residue was diluted with EtOAc. The organic layer was washed twice with saturated aqueous NH_4_Cl and brine, dried over MgSO_4_, and concentrated *in vacuo*. The residue was purified by column chromatography (silica gel, CHCl_3_ to CHCl_3_/EtOAc = 50:1) to afford compound **4** (52 mg, 0.122 mmol, 88%) as a white solid. ^1^H NMR (300 MHz, CDCl_3_): *δ* 8.56 (brs, 1H), 7.92 (s, 1H), 7.61 (dd, *J* = 5.2, 8.9 Hz, 2H), 7.20 (t, *J* = 8.9 Hz, 2H), 7.07 (d, *J* = 1.1 Hz, 1H), 5.23 (brs, 1H, NH), 3.75 (s, 3H), 3.25 (brs, 2H), 1.99 (brs, 1H), 1.84–1.68 (m, 4H), 1.33–1.15 (m, 4H), 1.08–0.96 (m, 2H). HRMS (ESI^+^) *m*/*z* Calcd. for C_22_H_25_FN_5_OS^+^ [M+H]^+^ 426.1758; Found 426.1750.

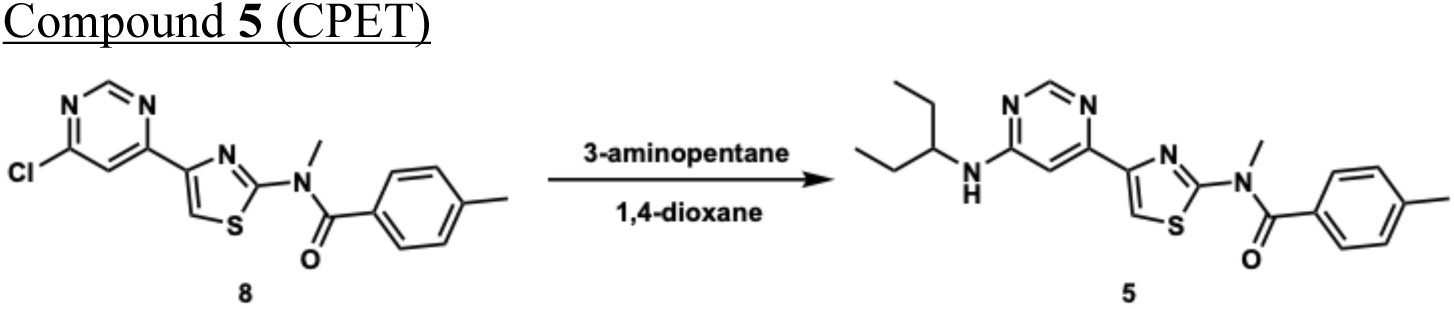

3-Aminopentane (2 ml, 17.2 mmol, 198 equiv.) was added to a mixture of compound **8** (30 mg, 0.087 mmol, 1 equiv.) in dry 1,4-dioxane (5 ml). The reaction mixture was stirred under N_2_ at 80–100 °C for 2 days. The mixture was concentrated *in vacuo* to remove the solvent and excess 3-aminopentane, and the residue was diluted with EtOAc. The organic layer was washed twice with saturated aqueous NH_4_Cl and brine, dried over Na_2_SO_4_, and concentrated *in vacuo*. The residue was purified by column chromatography (silica gel, *n*-hexane/EtOAc = 2:1 to 1:1) to afford compound **5** (29 mg, 0.073 mmol, 84%) as a white solid. ^1^H NMR (500 MHz, 1,1,2,2-tetrachloroethane-*d*_2_, 353 K): *δ* 8.57 (s, 1H), 8.17 (brs, 1H), 7.51 (d, *J* = 8.1 Hz, 2H), 7.34 (d, *J* = 8.1 Hz, 2H), 7.14 (s, 1H), 5.27 (brs, 1H, NH), 3.84 (brm, 1H), 3.78 (s, 3H), 2.48 (s, 3H), 1.78–1.56 (m, 4H), 1.02 (t, *J* = 7.4 Hz, 6H). HRMS (ESI^+^) *m*/*z* Calcd. for C_21_H_25_N_5_OS^+^ [M+H]^+^ 396.1853; Found 396.1852.

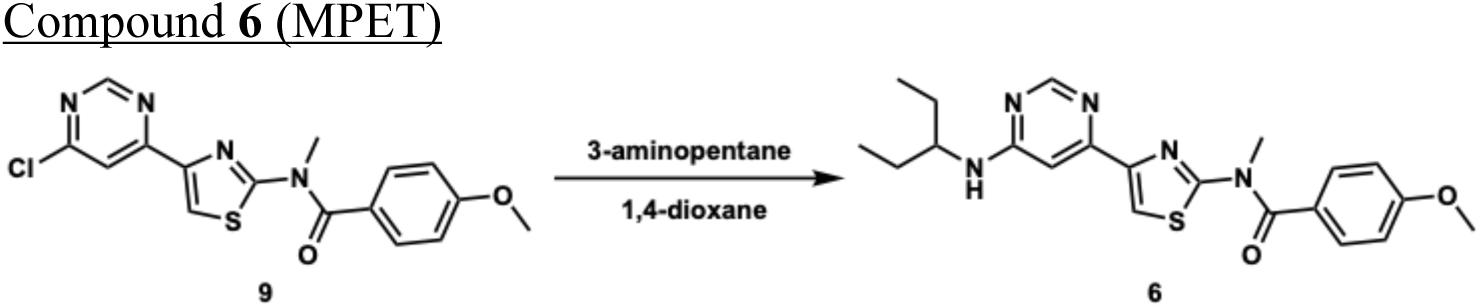

<u>3</u>-Aminopentane (1.1 ml, 9.46 mmol, 68 equiv.) was added to a mixture of compound **9** (50 mg, 0.139 mmol, 1 equiv.) in dry 1,4-dioxane (5 ml). The reaction mixture was stirred under N_2_ at 80 °C for 24 h. The mixture was concentrated *in vacuo* to remove the solvent and excess 3-aminopentane, and the residue was diluted with EtOAc. The organic layer was washed with saturated aqueous NH_4_Cl, water, and brine, dried over Na_2_SO_4_, and concentrated *in vacuo*. The residue was purified by column chromatography (silica gel, *n*-hexane/EtOAc = 4:1 to 1:2) to afford compound **6** (28 mg, 0.068 mmol, 49%) as a white solid. ^1^H NMR (500 MHz, 1,1,2,2-tetrachloroethane-*d*_2_, 353 K): *δ* 8.56 (d, *J* = 0.9 Hz, 1H), 7.94 (s, 1H), 7.60 (d, *J* = 8.8 Hz, 2H), 7.10 (d, *J* = 0.9 Hz, 1H), 7.04 (d, *J* = 8.8 Hz, 2H), 4.79 (brd, *J* = 8.7 Hz, 1H, NH), 3.92 (s, 3H), 3.82 (brm, 1H), 3.81 (s, 3H), 1.76–1.54 (m, 4H), 1.01 (t, *J* = 7.4 Hz, 6H). HRMS (ESI^+^) *m*/*z* Calcd. for C_21_H_25_N_5_O_2_S^+^ [M+H]^+^ 412.1802; Found 412.1808.

### Synthesis of the precursor of [^11^C]MPET

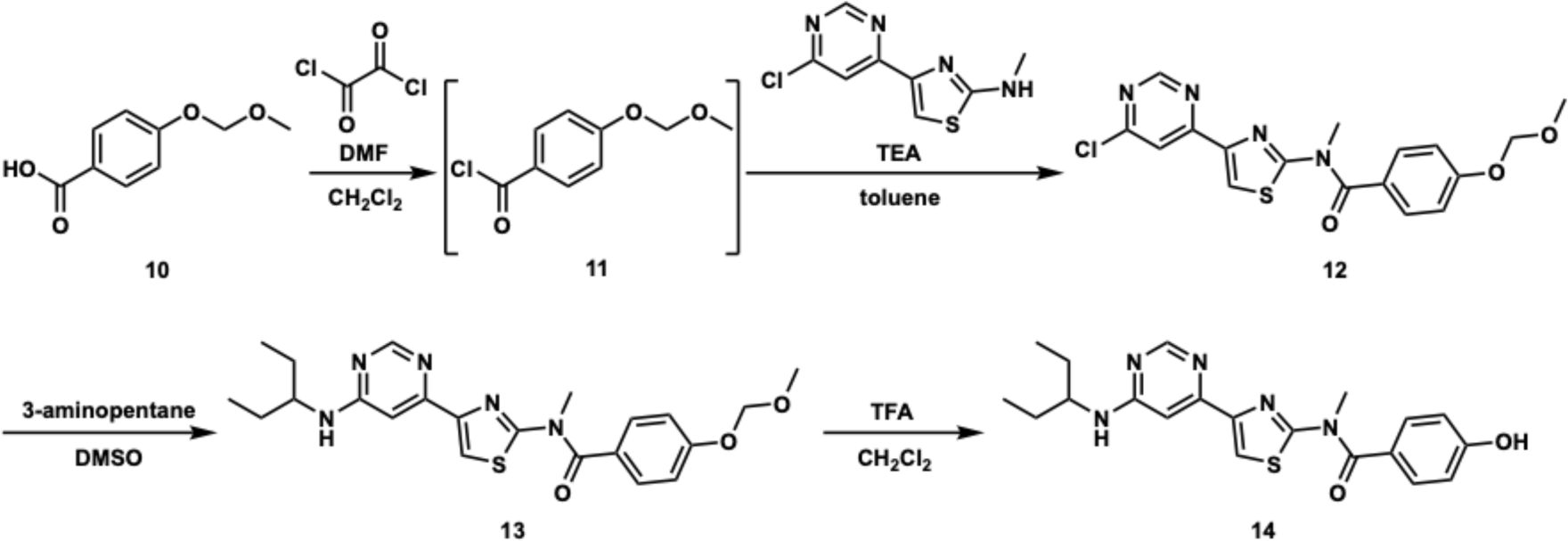

4-(6-Chloro-4-pyrimidinyl)-N-methyl-2-thiazolamine was synthesized according to a modified literature procedure (ref. S1). Compound **10** was synthesized according to a modified literature procedure (ref. S4).

#### Compound 12

To a solution of compound **10** (78 mg, 0.428 mmol, 1 equiv.) in dry dichloromethane (3 ml), dry DMF (3 µl, 0.039 mmol, 0.09 equiv.) and oxalyl chloride (110 µl, 1.28 mmol, 3 equiv.) were added under N_2_ in an ice bath. The reaction mixture was stirred at room temperature for 6 h and then concentrated *in vacuo*. The crude intermediate **11** was treated with 4-(6-chloro-4-pyrimidinyl)-N-methyl-2-thiazolamine (88 mg, 0.388 mmol, 0.9 equiv.), triethylamine (160 µl, 1.15 mmol, 2.7 equiv.) and dry toluene (3 ml) under N_2_. The reaction mixture was stirred at 100 °C overnight and concentrated *in vacuo*, and the residue was diluted with CHCl_3_. The organic layer was washed twice with saturated aqueous NaHCO_3_ and brine, dried over Na_2_SO_4_, and concentrated *in vacuo*. The residue was purified by column chromatography (silica gel, CHCl_3_/EtOAc = 100:1 to 9:1) to afford compound **12** (68 mg, 0.174 mmol, 41%) as a white solid. ^1^H NMR (300 MHz, CDCl_3_): *δ* 8.95 (d, *J* = 1.1 Hz, 1H), 8.10 (s, 1H), 8.06 (d, *J* = 1.1 Hz, 1H), 7.58 (d, *J* = 8.9 Hz, 2H), 7.15 (d, *J* = 8.9 Hz, 1H), 5.26 (s, 2H), 3.80 (s, 3H), 3.52 (s, 3H).

#### Compound 13

To a solution of compound **12** (68 mg, 0.174 mmol, 1 equiv.) in dry DMSO (3 ml), 3-aminopentane (101 µl, 0.867 mmol, 5 equiv.) was added under N_2_. The reaction mixture was stirred at 90 °C overnight. The mixture was diluted with EtOAc, and washed twice with saturated aqueous NaHCO_3_ and brine. The aqueous layer was extracted twice with EtOAc. The combined organic layers were dried over Na_2_SO_4_, and concentrated *in vacuo*. The residue was purified by column chromatography (silica gel, CHCl_3_/EtOAc = 3:1 containing 1% triethylamine) to afford compound **13** (44 mg, 0.100 mmol, 57%) as a white solid. ^1^H NMR (500 MHz, 1,1,2,2-tetrachloroethane-*d*_2_, 353 K): *δ* 8.56 (s, 1H), 7.95 (s, 1H), 7.59 (d, *J* = 8.7 Hz, 2H), 7.17 (d, *J* = 8.7 Hz, 2H), 7.10 (s, 1H), 5.27 (s, 2H), 4.78 (brd, *J* = 8.7 Hz, 1H, NH), 3.81 (brm, 4H), 3.55 (s, 3H), 1.76–1.55 (m, 4H), 1.01 (t, *J* = 7.4 Hz, 6H).

#### Compound 14

To a solution of compound **13** (22 mg, 0.050 mmol, 1 equiv.) in dry dichloromethane (3 ml), trifluoroacetic acid (500 µl, 6.53 mmol, 131 equiv.) was added under N_2_. The reaction mixture was stirred at room temperature for 2 h, and then concentrated *in vacuo*. The residue was purified by column chromatography (silica gel, CHCl_3_/MeOH = 9:1 containing 1% triethylamine) to afford compound **14** (16 mg, 0.040 mmol, 80%) as a white solid. ^1^H NMR (500 MHz, 1,1,2,2-tetrachloroethane-*d*_2_, 353 K): *δ* 8.57 (s, 1H), 7.95 (s, 1H), 7.55 (d, *J* = 8.5 Hz, 2H), 7.11 (s, 1H), 6.97 (t, *J* = 8.5 Hz, 2H), 4.81 (brd, *J* = 8.0 Hz, 1H, NH), 3.80 (brm, 4H), 1.76–1.54 (m, 4H), 1.01 (t, *J* = 7.4 Hz, 6H). HRMS (ESI^+^) *m*/*z* Calcd. for C_20_H_23_N_5_O_2_S^+^ [M+H]^+^ 398.1645; Found 398.1654.

## References

1. Riedel, G., Platt, B. & Micheau, J. Glutamate receptor function in learning and memory. Behav. Brain Res. 140, 1–47 (2003).

2. Lovinger, D. M. Neurotransmitter roles in synaptic modulation, plasticity and learning in the dorsal striatum. Neuropharmacology 58, 951–961 (2010).

3. Niswender, C. M. & Conn, P. J. Metabotropic glutamate receptors: physiology, pharmacology, and disease. Anuu. Rev. Pharmacol. Toxicol. 50, 295–322 (2010).

4. Martel, J. C. & McArthur, S. G. Dopamine receptor subtypes, physiology and pharmacology: new ligands and concepts in schizophrenia. Front. Pharmacol. 11, 1003 (2020).

5. Wess, J., Eglen, R. M. & Gautam. D. Muscarinic acetylcholine receptors: mutant mice provide new insights for drug development. Nat. Rev. Drug Discov. 6, 721–733 (2007).

6. Skarnes, W. C. et al. A conditional knockout resource for the genome-wide study of mouse gene function. Nature 474, 337–342 (2011).

7. Schöneberg, T. & Liebscher, I. Mutations in G protein-coupled receptors: mechanisms, pathophysiology and potential therapeutic approaches. Pharmacol. Rev. 73, 89–119 (2011).

8. El-Brolosy, M. A. & Stainier, D. Y. R. Genetic compensation: a phenomenon in search of mechanisms. PLoS Genet. 13, e1006780 (2017).

9. Aiba, A. & Nakao, H. Conditional mutant mice using tetracycline-controlled gene expression system in the brain. Neurosci. Res. 58, 113–117 (2007).

10. Snyder, S. H. Drug and neurotransmitter receptors in the brain. Science 224, 22–31 (1984).

11. Peters, J.-U. Polypharmacology – foe or friend? J. Med. Chem. 56, 8955–8971 (2013).

12. Sutherland, J. J., Yonchev, D., Fekete, A. & Urban, L. A preclinical secondary pharmacology resource illuminates target-adverse drug reaction associations of marketed drugs. Nat. Commun. 14, 4323 (2023).

13. Campbell, E. J. & Marchant, N. J. The use of chemogenetics in behavioural neuroscience: receptor variants, targeting approaches and caveats. Br. J. Pharmacol. 175, 994–1003 (2018).

14. Miura, Y., Senoo, A., Doura, T. & Kiyonaka, S. Chemogenetics of cell surface receptors: beyond genetic and pharmacological approaches. RSC Chem. Biol. 3, 269–287 (2022).

15. Roth, B. L. DREADDs for neuroscientists. Neuron 89, 683–694 (2016).

16. Ryo, Y., Miyawaki, A., Furuichi, T. & Mikoshiba, K. Expression of the metabotropic glutamate receptor mGluR1α and the ionotropic glutamate receptor GluR1 in the brain during the postnatal development of normal mouse and in the cerebellum from mutant mice. J. Neurosci. Res. 36, 19–32 (1993).

17. Nakao, H., Nakao, K., Kano, M. & Aiba, A. Metabotropic glutamate receptor subtype-1 is essential for motor coordination in the adult cerebellum. Neurosci. Res. 57, 538–543 (2007).

18. Ichise, T. et al. mGluR1 in cerebellar Purkinje cells essential for long-term depression, synapse elimination, and motor coordination. Science 288, 1832–1835 (2000).

19. Satoh, A. et al. Discovery and in vitro and in vivo profiles of 4-fluoro-*N*-[4-[6-(isopropylamino)pyrimidin-4-yl]-1,3-thiazol-2-yl]-*N*-methylbenzamide as novel class of an orally active metabotropic glutamate receptor 1 (mGluR1) antagonist. Bioorg. Med. Chem. Lett. 19, 5464–5468 (2009).

20. Wu, H. et al. Structure of a class C GPCR metabotropic glutamate receptor 1 bound to an allosteric modulator. Science 344, 58–64 (2014).

21. Ojima, K. et al. Coordination chemogenetics for activation of GPCR-type glutamate receptors in brain tissue. Nat. Commun. 13, 3167 (2022).

22. Zhou, Q. et al. Common activation mechanism of class A GPCRs. eLife 8, e50279 (2019).

23. Yamasaki, T. et al. Imaging for metabotropic glutamate receptor subtype 1 in rat and monkey brains using PET with [^18^F]FITM. Eur. J. Nucl. Med. Mol. Imaging 39, 632–641 (2012).

24. Fujinaga, M. et al. Development of *N*-[4-[6-(isopropylamino)pyrimidin-4-yl]-1,3-thiazol-2-yl]-*N*-methyl-4-[^11^C]methylbenzamide for positron emission tomography imaging of metabotropic glutamate 1 receptor in monkey brain. J. Med. Chem. 55, 11042–11051 (2012).

25. Fujinaga, M. et al. Synthesis and evaluation of novel radioligands for positron emission tomography imaging of metabotropic glutamate receptor subtype 1 (mGluR1) in rodent brain. J. Med. Chem. 55, 2342–2352 (2012).

26. Santoro, E. M. F. et al. Purkinje cell models: past, present and future. Front. Comput. Neurosci. 18, 1426653 (2024).

27. Linden, D. J. & Connor, J. A. Cellular mechanisms of long-term depression in the cerebellum. Curr. Opin. Neurobiol. 3, 401–406 (1993).

28. Aiba, A. et al. Deficient cerebellar long-term depression and impaired motor learning in mGluR1 mutant mice. Cell 79, 377–388 (1994).

29. van Alphen, A. M., Stahl, J. S. & De Zeeuw, C. I. The dynamic characteristics of the mouse horizontal vestibulo-ocular and optokinetic response. Brain Res. 890, 296–305 (2001).

30. Iwashita, M., Kanai, R., Funabiki, K., Matsuda, K. & Hirano, T. Dynamic properties, interactions and adaptive modifications of vestibulo-ocular reflex and optokinetic response in mice. Neurosci. Res. 39, 299–311 (2001).

31. Shutoh, F. et al. Loss of adaptability of horizontal optokinetic response eye movements in mGluR1 knockout mice. Neurosci. Res. 42, 141–145 (2002).

32. Shiotsuki, H. et al. A rotarod test for evaluation of motor skill learning. J. Neurosci. Methods 189, 180–185 (2010).

33. Kakegawa, W. et al. D-Serine regulates cerebellar LTD and motor coordination through the δ2 glutamate receptor. Nat. Neurosci. 14, 603–611 (2011).

34. Armbruster, B. N., Li, X., Pausch, M. H., Herlitze, S. & Roth, B. L. Evolving the lock to fit the key to create a family of G protein-coupled receptors potently activated by an inert ligand. Proc. Natl. Acad. Sci. USA 104, 5163–5168 (2007).

35. Nagai, Y. et al. Deschloroclozapine, a potent and selective chemogenetic actuator enables rapid neuronal and behavioral modulations in mice and monkeys. Nat. Neurosci. 23, 1157–1167 (2020).

36. Kalogriopoulos, N. A. et al. Synthetic GPCRs for programmable sensing and control of cell behaviour. Nature 637, 230–239 (2025).

37. Magnus, C. J. et al. Chemical and genetic engineering of selective ion channel–ligand interactions. Science 333, 1292–1296 (2011).

38. Magnus, C. J. et al. Ultrapotent chemogenetics for research and potential clinical applications. Science 364, eaav5282 (2019).

39. Shields, B. C. et al. Deconstructing behavioral neuropharmacology with cellular specificity. Science 356, eaaj2161 (2017).

40. Shields, B. C., et al. DART.2: bidirectional synaptic pharmacology with thousandfold cellular specificity. Nat. Methods 21, 1288–1297 (2024).

41. Los, G. V. et al. HaloTag: a novel protein labeling technology for cell imaging and protein analysis. ACS Chem. Biol. 3, 373–382 (2008).

42. Donthamsetti, P. C. et al. Genetically targeted optical control of an endogenous G protein-coupled receptor. J. Am. Chem. Soc. 141, 11522–11530 (2019).

43. Donthamsetti, P. et al. Cell specific photoswitchable agonists for reversible control of endogenous dopamine receptors. Nat. Commun. 12, 4775 (2021).

44. Kobauri, P., Dekker, F. J., Szymanski, W. & Feringa, B. L. Rational design in photopharmacology with molecular photoswitches. Angew. Chem. Int. Ed. 62, e202300681 (2023).

45. Shingles, G. et al. A chemogenetic approach for temporal and cell-specific activation of endogenous GPCRs in vivo. Proc. Natl. Acad. Sci. USA 123, e2501228123 (2026).

46. Kiyonaka, S. et al. Allosteric activation of membrane-bound glutamate receptors using coordination chemistry within living cells. Nat. Chem. 8, 958–967 (2016).

47. Shcherbakova, D. M. & Verkhusha, V. V. Near-infrared fluorescent proteins for multicolor *in vivo* imaging. Nat. Methods 10, 751–754 (2013).

48. Söding, J. Protein homology detection by HMM–HMM comparison. Bioinformatics 21, 951–960 (2005).

49. Šali, A. & Blundell, T. L. Comparative protein modelling by satisfaction of spatial restraints. J. Mol. Biol. 234, 779–815 (1993).

50. Tian, C. et al. ff19SB: Amino-acid-specific protein backbone parameters trained against quantum mechanics energy surfaces in solution. J. Chem. Theory Comput. 16, 528–552 (2020).

51. Wang, J., Wolf R. M., Caldwell, J. W., Kollman, P. A. & Case, D. A. Development and testing of a general amber force field. J. Comput. Chem. 25, 1157–1174 (2004).

52. Dickson, C. J., Walker, R. C. & Gould, I. R. Lipid21: Complex lipid membrane simulations with AMBER. J. Chem. Theory Comput. 18, 1726–1736 (2022).

53. Chamorro, V. C., Jungwirth, P. & Martinez-Seara, H. Building water models compatible with charge scaling molecular dynamics. J. Phys. Chem. Lett. 15, 2922–2928 (2024).

54. Li, P., Song, L. F. & Merz, K. M. Jr. Systematic parameterization of monovalent ions employing the nonbonded model. J. Chem. Theory Comput. 11, 1645–1657 (2015).

55. Åqvist, J., Wennerström, P., Nervall, M., Bjelic, S. & Brandsdal, B. O. Molecular dynamics simulations of water and biomolecules with a Monte Carlo constant pressure algorithm. Chem. Phys. Lett. 384, 288–294 (2004).

56. Suzuki, K. et al. Computer-controlled large scale production of high specific activity [^11^C]RO 15-1788 for PET studies of benzodiazepine receptors. Int. J. Appl. Radiat. Isot. 36, 971–976 (1985).

57. Bertoglio, D. et al. In vitro and in vivo assessment of suitable reference region and kinetic modelling for the mGluR1 radioligand [^11^C]ITDM in mice. Mol. Imaging Biol. 22, 854–863 (2020).

58. Obokata, N. et al. Synthesis and preclinical evalution of [^11^C]MTP38 as a novel PET ligand for phosphodiesterase 7 in the brain. Eur. J. Nucl. Med. Mol. Imaging 48, 3101–3112 (2021).

59. Lassen, N. A. et al. Benzodiazepine receptor quantification in vivo in humans using [^11^C]flumazenil and PET: application of the steady-state principle. J. Cereb. Blood Flow Metab. 15, 152–165 (1995).

60. Innis, R. B. et al. Consensus nomenclature for in vivo imaging of reversibly binding radioligands. J. Cereb. Blood Flow Metab. 27, 1533–1539 (2007).

61. Kakegawa, W. et al. Optogenetic control of synaptic AMPA receptor endocytosis reveals roles of LTD in motor learning. Neuron 99, 985–998.e6 (2018).

62. Nagao, S. Eye velocity is not the major factor that determines mossy fiber responses of rabbit floccular Purkinje cells to head and screen oscillation. Exp. Brain Res. 80, 221–224 (1990).

## Supplementary References

S1. Satoh, A. et al. Discovery and in vitro and in vivo profiles of 4-fluoro-N-[4-[6-(isopropylamino)pyrimidin-4-yl]-1,3-thiazol-2-yl]-N-methylbenzamide as novel class of an orally active metabotropic glutamate receptor 1 (mGluR1) antagonist. Bioorg. Med. Chem. Lett. 19, 5464–5468 (2009).

S2. Fujinaga, M. et al. Development of N-[4-[6-(isopropylamino)pyrimidin-4-yl]-1,3-thiazol-2-yl]-N-methyl-4-[11C]methylbenzamide for positron emission tomography imaging of metabotropic glutamate 1 receptor in monkey brain. J. Med. Chem. 55, 11042–11051 (2012).

S3. Fujinaga, M., et al. Synthesis and evaluation of novel radioligands for positron emission tomography imaging of metabotropic glutamate receptor subtype 1 (mGluR1) in rodent brain. J. Med. Chem. 55, 2342–2352 (2012).

S4. Lampe, J. W., et al. Synthesis and protein kinase inhibitory activity of balanol analogues with modified benzophenone subunits. J. Med. Chem. 45, 2624–2643 (2002).

